# Extracellular electron transfer-dependent anaerobic oxidation of ammonium by anammox bacteria

**DOI:** 10.1101/855817

**Authors:** Dario R. Shaw, Muhammad Ali, Krishna P. Katuri, Jeffrey A. Gralnick, Joachim Reimann, Rob Mesman, Laura van Niftrik, Mike S. M. Jetten, Pascal E. Saikaly

**Affiliations:** Water Desalination and Reuse Center (WDRC), Biological and Environmental Science & Engineering (BESE) Division, King Abdullah University of Science and Technology (KAUST), Thuwal 23955-6900, Saudi Arabia; BioTechnology Institute and Department of Plant and Microbial Biology, University of Minnesota, Twin Cities, St. Paul, Minnesota 55108, United States; Department of Microbiology, Institute for Water and Wetland Research (IWWR), Faculty of Science, Radboud University, Heyendaalseweg 135, 6525 AJ Nijmegen, the Netherlands

## Abstract

Anaerobic ammonium oxidation (anammox) by anammox bacteria contributes significantly to the global nitrogen cycle, and plays a major role in sustainable wastewater treatment. Anammox bacteria convert ammonium (NH_4_^+^) to dinitrogen gas (N_2_) using nitrite (NO_2_^−^) or nitric oxide (NO) as the electron acceptor. In the absence of NO_2_^−^ or NO, anammox bacteria can couple formate oxidation to the reduction of metal oxides such as Fe(III) or Mn(IV). Their genomes contain homologs of *Geobacter* and *Shewanella* cytochromes *involved in extracellular* electron transfer (EET). However, it is still unknown whether anammox bacteria have EET capability and can couple the oxidation of NH_4_^+^ with transfer of electrons to carbon-based insoluble extracellular electron acceptors. Here we show using complementary approaches that in the absence of NO_2_^−^, freshwater and marine anammox bacteria couple the oxidation of NH_4_^+^ with transfer of electrons to carbon-based insoluble extracellular electron acceptors such as graphene oxide (GO) or electrodes poised at a certain potential in microbial electrolysis cells (MECs). Metagenomics, fluorescence *in-situ* hybridization and electrochemical analyses coupled with MEC performance confirmed that anammox electrode biofilms were responsible for current generation through EET-dependent oxidation of NH_4_^+^. ^15^N-labelling experiments revealed the molecular mechanism of the EET-dependent anammox process. NH_4_^+^ was oxidized to N_2_ via hydroxylamine (NH_2_OH) as intermediate when electrode was the terminal electron acceptor. Comparative transcriptomics analysis supported isotope labelling experiments and revealed an alternative pathway for NH_4_^+^ oxidation coupled to EET when electrode is used as electron acceptor compared to NO_2_^−^as electron acceptor. To our knowledge, our results provide the first experimental evidence that marine and freshwater anammox bacteria can couple NH_4_^+^ oxidation with EET, which is a significant finding, and challenges our perception of a key player of anaerobic oxidation of NH_4_^+^ in natural environments and engineered systems.

---

Anaerobic ammonium oxidation (anammox) by anammox bacteria contributes up to 50% of N_2_ emitted into Earth’s atmosphere from the oceans (*1, 2*). Also, anammox bacteria has been extensively investigated for energy-efficient removal of NH_4_^+^ from wastewater (*3*). Initially, anammox bacteria were assumed to be restricted to NH_4_^+^ as electron donor and NO_2_^−^ or NO as electron acceptor (*4, 5*). More than a decade ago, preliminary experiments showed that *Kuenenia stuttgartiensis* and *Scalindua* could couple the oxidation of formate to the reduction of insoluble extracellular electron acceptors such as Fe(III) or Mn(IV) oxides (*6, 7*). However, the mechanism of how anammox bacteria reduce insoluble extracellular electron acceptors has remained unexplored to date. Also, growth or electrochemical activity was not quantified in these experiments. Further, these experiments could not discriminate between Fe(III) oxide reduction for nutritional acquisition (i.e., via siderophores) versus respiration through extracellular electron transfer (EET) (*8*). Therefore, with these preliminary experiments it could not be determined if anammox bacteria have EET capability or not.

Although preliminary work showed that *K. stuttgartiensis* could not reduce Mn(IV) or Fe(III) with NH_4_^+^ as electron donor (*6*), the possibility of anammox bacteria to oxidize NH_4_^+^ coupled to EET to other insoluble extracellular electron acceptors cannot be ruled out. In fact EET (and set of genes involved with EET) is not uniformly applied to all insoluble extracellular electron acceptors; some electroactive bacteria are not able to transfer electrons to carbon-based insoluble extracellular electron acceptors such as electrodes in bioelectrochemical systems but could reduce metal oxides and vice versa (*9*). It is known for more than two decades that carbon-based high-molecular-weight organic materials, which are ubiquitous in terrestrial and aquatic environments and that are not involved in microbial metabolism (i.e., humic substances) can be used as external electron acceptor for the anaerobic oxidation of compounds (*10*). Also, it has been reported that anaerobic NH_4_^+^ oxidation linked to microbial reduction of natural organic matter fuels nitrogen loss in marine sediments (*11*). A literature survey of more than 100 EET-capable species indicated that there are many ecological niches for microorganisms able to perform EET (*12*). This resonates with a recent finding where *Listeria monocytogenes*, a host-associated pathogen and fermentative Gram-positive bacterium, was able to respire through a flavin-based EET process and behaved as an electrochemically active microorganism (i.e., able to transfer electrons from oxidized fuel (substrate) to a working electrode via EET process) (*13*). Further it was reported that anammox bacteria seem to have homologs of *Geobacter* and *Shewanella* multi-heme cytochromes that are responsible *for* EET (*14*). These observations stimulated us to investigate whether anammox bacteria can couple NH_4_^+^ oxidation with EET to carbon-based insoluble extracellular electron acceptor and can behave as electrochemically active bacteria.

## Ammonium oxidation coupled with EET

To evaluate if anammox bacteria possess EET capability, we first tested whether enriched cultures of three phylogenetically and physiologically distant anammox species can couple the oxidation of NH_4_^+^ with the reduction of insoluble extracellular electron acceptor. Cultures of *Ca*. Brocadia (freshwater anammox species) and *Ca*. Scalindua (marine anammox species) were enriched and grown as planktonic cells in membrane bioreactors (Fig. S1A) (*15*). Fluorescence in situ hybridization (FISH) showed that the anammox bacteria constituted >95% of the bioreactor’s community (Fig. S1B-G). Also, a previously enriched *K. stuttgartiensis* (freshwater anammox species) culture was used (*4*). The anammox cells were incubated anoxically for 216 hours in the presence of ^15^NH_4_^+^ (4 mM) and graphene oxide (GO) as a proxy for insoluble electron acceptor. No NO_2_^−^ or NO_3_^−^ were added to the incubations. GO particles are bigger than bacterial cells and cannot be internalized, and thus GO can only be reduced by EET (*16*). Indeed, GO was reduced by anammox bacteria as shown by the formation of suspended reduced GO (rGO), which is black in color and insoluble (Fig. 1A) (*16*). In contrast, abiotic controls did not form insoluble black precipitates. Reduction of GO to rGO by anammox bacteria was further confirmed by Raman spectroscopy, where the formation of the characteristic 2D and D+D′ peaks of rGO (*17*) were detected in the vials with anammox cells (Fig. 1B), whereas no peaks were detected in the abiotic control. Further, isotope analysis of the produced N_2_ gas showed that anammox cells were capable of ^30^N_2_ formation (Fig. 1C). In contrast, ^29^N_2_ production was not significant in any of the tested anammox species or controls, suggesting that unlabeled NO_2_^−^ or NO_3_^−^ were not involved. The production of ^30^N_2_ indicated that the anammox cultures use a different mechanism for NH_4_^+^ oxidation in the presence of an insoluble extracellular electron acceptor (further explained below). Gas production was not observed in the abiotic control (Fig. 1C). To determine if anammox bacteria are still dominant after incubation with GO, we extracted and sequenced total DNA from the *Brocadia* and *Scalindua* vials at the end of the experiment. Differential coverage showed that the metagenomes were dominated by anammox bacteria (Fig. S2A and C). Also, no known EET-capable bacteria were detected in the metagenomes. Taken together, these results support that anammox bacteria have EET capability.

**Figure 1.**
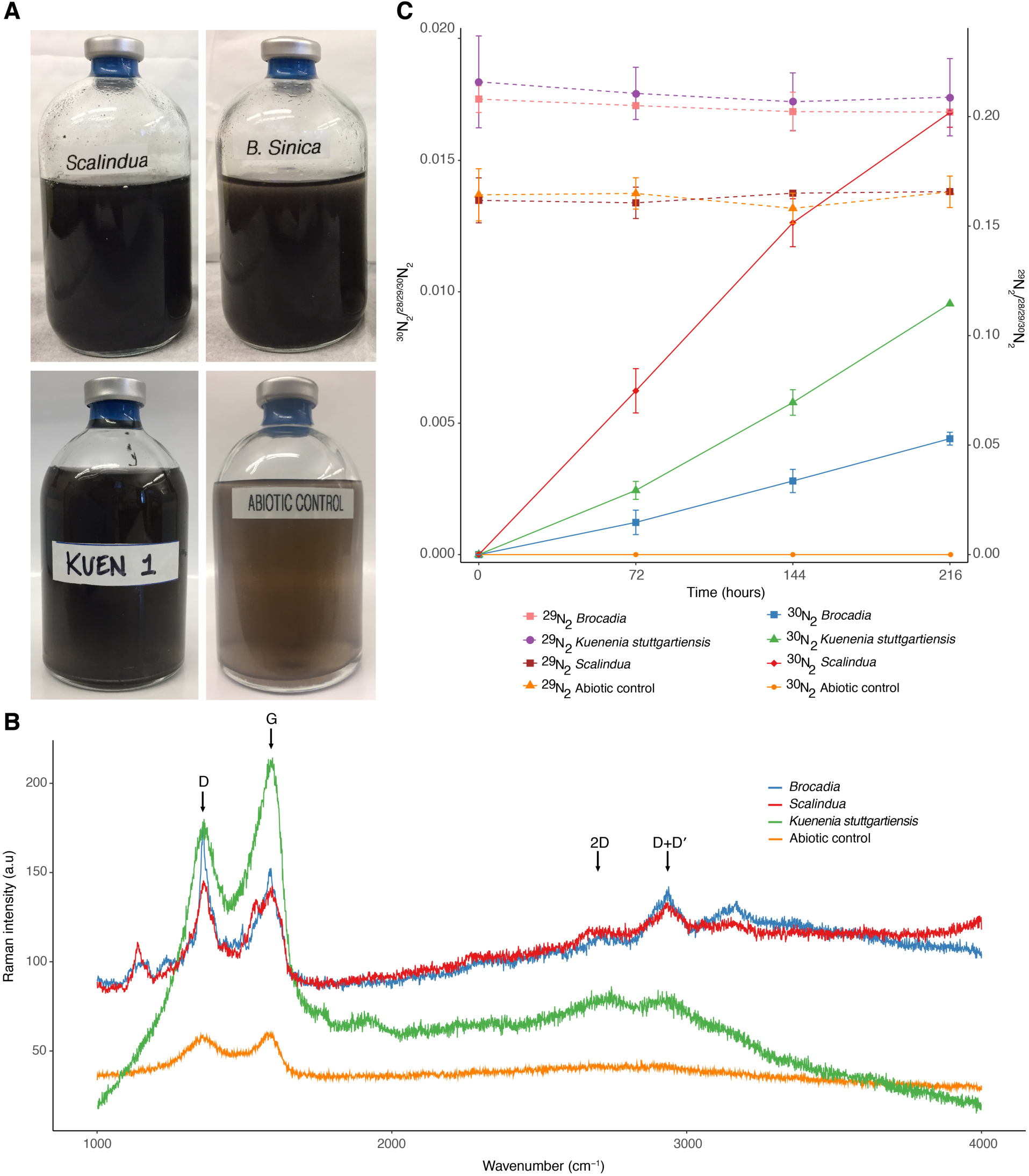
Different anammox bacteria can perform EET by coupling the oxidation of NH_4_^+^ with the reduction of GO. (**A**) Photographs of serum vials after 216 hours of incubation with different species of anammox bacteria, ^15^NH_4_^+^ and GO. The presence of black precipitates indicates the formation of reduced GO (rGO). No obvious change in color was observed in the abiotic control vials after the same period of incubation with ^15^NH_4_^+^ and GO. (**B**) Raman spectra of the vials after 216 hours of incubation. Peaks in bands of 2D and D + D′ located at ∼2700 and ∼2900 cm^-1^, respectively, indicate the formation of rGO. (**C**) ^30^N_2_ production by different anammox bacteria from ^15^NH_4_^+^ and GO as the sole electron acceptor. Anammox cells were incubated with 4 mM ^15^NH_4_^+^ and GO to a final concentration of 200 mg L^-1^. There was no ^29^N_2_ formation throughout the experiment. NO and N_2_O were not detected throughout the experiment. Results from triplicate serum vial experiments are represented as mean ± SD.

## Electroactivity of anammox bacteria

Electrochemical techniques provide a powerful tool to evaluate EET, where electrodes substitute for the insoluble minerals as the terminal electron acceptor (*13*). Compared to metal oxides, the use of electrodes as the terminal electron acceptor allow us to quantify the number of externalized electrons per mol of NH_4_^+^ oxidized. Also, since the electrode is only used for bacterial respiration, then we can better assess EET activity compared to metal oxides, where we cannot differentiate between metal oxide reduction for nutritional acquisition from respiration through EET activity. Therefore, we tested if anammox bacteria interact with electrodes via EET and use them as the sole electron acceptor in MEC. Single-chamber MEC operated at eight different set potentials (from –0.3 to 0.4 V vs Ag/AgCl) using multiple working electrodes (Fig. S1H) were initially operated under abiotic conditions with the addition of NH_4_^+^ only. No current and NH_4_^+^ removal were observed in any of the abiotic controls. Subsequently, the *Ca*. Brocadia culture was inoculated into the MEC and operated under optimal conditions for anammox (i.e., addition of NH_4_^+^ and NO_2_^−^). Under this scenario, NH_4_^+^ and NO_2_^−^ were completely removed from the medium without any current generation (Fig. 2A). Stoichiometric ratios of consumed NO_2_^−^ to consumed NH_4_^+^ (ΔNO_2_^−^/ΔNH_4_^+^) and produced NO_3_^−^ to consumed NH_4_^+^ (ΔNO_3_^−^/ΔNH_4_^+^) were in the range of 1.0–1.3 and 0.12–0.18, respectively, which are close to the theoretical ratios of the anammox reaction (*18*). These ratios indicated that anammox bacteria were responsible for NH_4_^+^ removal in the MEC. Subsequently, NO_2_^−^ was gradually decreased to 0 mM leaving the electrodes as the sole electron acceptor. When the exogenous electron acceptor (i.e., NO_2_^−^) was completely removed from the feed, anammox cells began to form a biofilm on the surface of the electrodes (Fig. S1I) and current generation coupled to NH_4_^+^ oxidation was observed in the absence of NO_2_^−^ (Fig. 2A). Further, NO_2_^−^ and NO_3_^−^ were below the detection limit at all time points when the working electrode was used as the sole electron acceptor. The magnitude of current generation was proportional to the NH_4_^+^ concentration (Fig. 2A) and maximum current density was observed at set potential of 0.4 V vs Ag/AgCl. There was no visible biofilm growth and current generation at set potentials ≤ 0 V vs Ag/AgCl. To confirm that the electrode-dependent anaerobic oxidation of NH_4_^+^ was catalyzed by anammox bacteria, additional control experiments were conducted in chronological order in the MEC. The presence of ATU, a compound that selectively inhibits aerobic NH_3_ oxidation by ammonia monooxygenase (AMO) in ammonia oxidizing bacteria (AOB), ammonia oxidizing archaea (AOA) and Comammox (*19*), did not result in an inhibitory effect on NH_4_^+^ removal and current generation (Fig. 2A). NH_4_^+^ was not oxidized when the MEC was operated in open circuit voltage mode (OCV; electrode is not used as electron acceptor) (Fig. 2B), strongly suggesting an electrode-dependent NH_4_^+^ oxidation and that trace amounts of O_2_ if present, are not responsible for NH_4_^+^ oxidation. Addition of NO_2_^−^ resulted in an immediate drop in current density with simultaneous removal of NH_4_^+^ and NO_2_^−^ and formation of NO_3_^−^, in the expected stoichiometry (*18*) (Fig. 2C). Repeated addition of NO_2_^−^ resulted in the complete abolishment of current generation, indicating that anammox bacteria were solely responsible for current production in the absence of an exogenous electron acceptor. Absence of NH_4_^+^ from the feed resulted in no current generation, and current was immediately resumed when NH_4_^+^ was added again to the feed (Fig. 2D), further supporting the role of anammox bacteria in current generation. These results also indicate that current generation was not catalyzed by electrochemically active heterotrophs, which might utilize organic carbon generated from endogenous decay processes. Autoclaving the MECs immediately stopped current generation and NH_4_^+^ removal (Fig. 2D) indicating that current generation was due a biotic reaction. Similar results were also obtained with MECs operated with *Ca*. Scalindua or *K. stuttgartiensis* cultures (Fig. S3A and B), suggesting that they are also electrochemically active and can oxidize NH_4_^+^ using working electrodes as the electron acceptor. Taken together these results provide strong evidence for electrode-dependent anaerobic oxidation of NH_4_^+^ by phylogenetically distant anammox bacteria.

**Figure 2.**
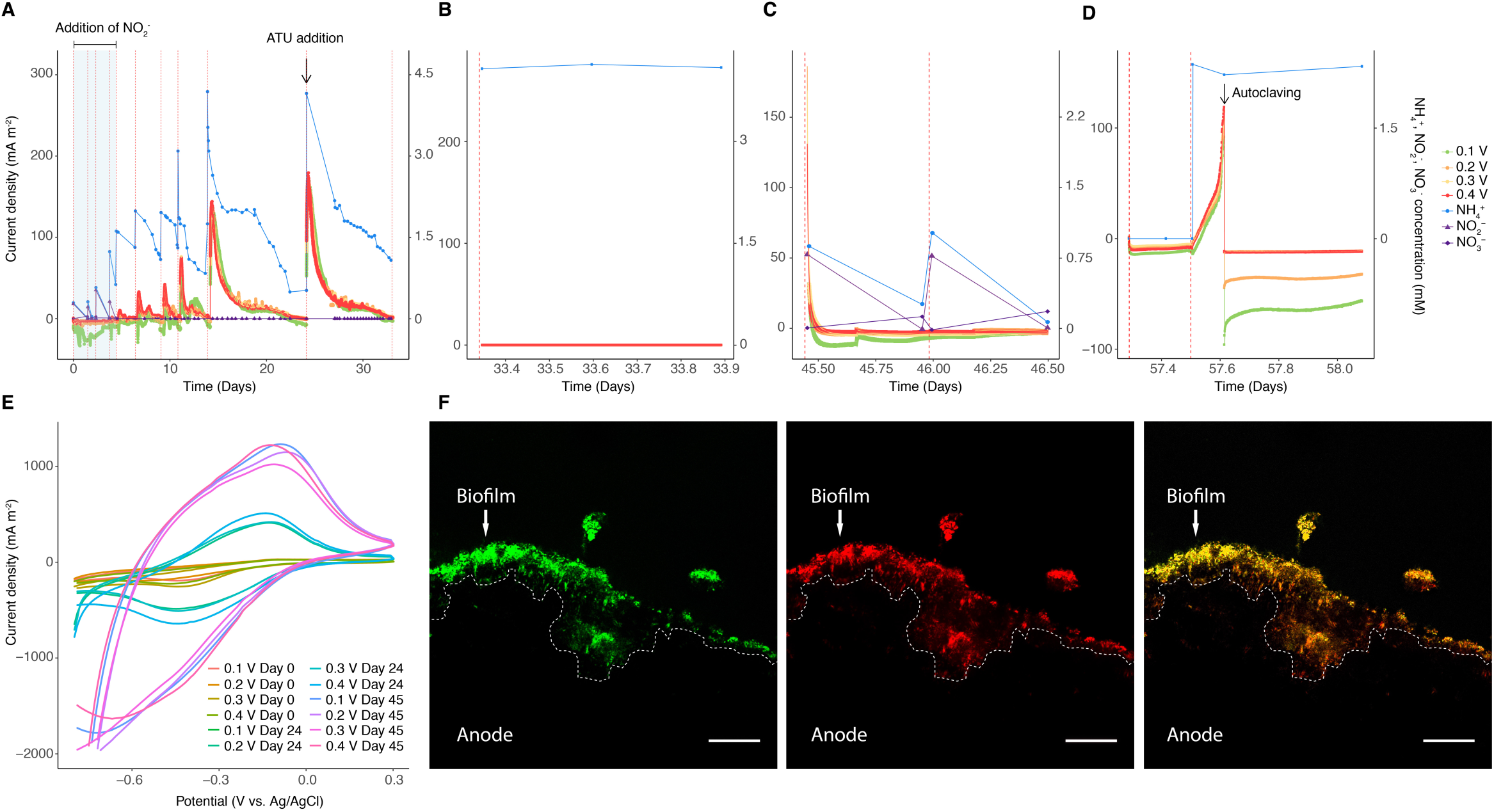
*Ca.* Brocadia is electrochemically active (i.e., able to release electrons from inside the cell to working electrode). (**A** to **D**) Ammonium oxidation coupled to current generation in chronoamperometry experiment conducted in single-chamber multiple working electrode MEC inoculated with *Ca.* Brocadia. (**A**) MEC operated initially under different set potentials with addition of nitrite, which is the preferred electron acceptor for ammonium oxidation by anammox bacteria, followed by operation with working electrodes as sole electron acceptors. The highlighted area in blue refers to the operation of MEC in the presence of nitrite. The black arrow indicates the addition of ATU, a compound that selectively inhibits nitrifiers. (**B**) MEC operated under open circuit voltage (OCV) mode. (**C**) MEC operated at different set potentials and with addition of nitrite. (**D**) MEC operated at different set potentials and without addition of ammonium and then with addition of ammonium followed by autoclaving. The black arrow in (**D**) indicates autoclaving followed by re-connecting of the MECs. Red dashed lines in (**A**), (**B**), (**C**) and (**D**) represent a change of batch. (**E**) Cyclic Voltammogram (1 mV s^-1^) of anammox biofilm grown on working electrodes (i.e., anodes) operated at different set potentials and growth periods following inoculation in MEC. (**F**) Confocal laser scanning microscopy images of a thin cross-section of the graphite rod anodes (0.4 V vs Ag/AgCl applied potential). The images are showing the *in-situ* spatial organization of all bacteria (green), anammox bacteria (red) and the merged micrograph (yellow). Fluorescence *in-situ* hybridization was performed with EUB I, II and III probes for all bacteria and Alexa647-labeled Amx820 probe for anammox bacteria. The dotted outline indicates the graphite rod anode surface. The white arrow indicates the biofilm. The scale bars represent 20 μm in length.

Cyclic voltammetry (CV) was used to correlate between current density and biofilm age, in cell-free filtrates (filtered reactor solution) and the developed biofilms at different time intervals. The anodes exhibited similar redox peaks with midpoint potentials (E_1/2_) of ∼200 mV vs Standard Hydrogen Electrode (SHE) for all three anammox species (Fig. 2E and Fig. S3C and D). No redox peaks were observed for the cell-free solution, indicating that soluble mediators are not involved in EET. Also, addition of exogenous riboflavin, which is a common soluble mediator involved in flavin-based EET process in gram-positive and gram-negative bacteria (*13, 20*), did not invoke changes in current density. Thus, the CV analysis corroborated that the electrode biofilms were responsible for current generation through direct EET mechanism.

The mole of electrons transferred to the electrode per mole of NH_4_^+^ oxidized to N_2_ (Table S1) was stoichiometrically close to equation 1 (Eq. 1). Also, electron balance calculations showed that coulombic efficiency (CE) was >80% for all NH_4_^+^ concentrations and anammox cultures tested in the experiments with electrodes as the sole electron acceptor (Table S1).

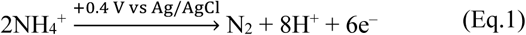

To determine if cathodic reaction (i.e., hydrogen evolution reaction) has an effect on electrode-dependent anaerobic NH_4_^+^ oxidation, additional experiments with *Ca*. Brocadia were conducted by operating single and double-chamber MECs in parallel (at 0.4 V vs Ag/AgCl applied potential). However, there was no significant difference in NH_4_^+^ oxidation and current production between the different reactor configurations (Fig. S4), suggesting no influence of cathodic reaction (i.e., H_2_ recycling) on the process. This was further supported by electron balance and CE calculations (Table S1). In addition, NH_4_^+^ oxidation and current production were not affected by the addition of Penicillin G (Fig. S4), a compound that has inhibitory effects in some heterotrophs but it does not have any observable short-term effects on anammox activity (*21, 22*). This further supports that current generation was not catalyzed by electrochemically active heterotrophs. Similar results were obtained with *Ca*. Scalindua and *K. stuttgartiensis* (data not shown).

Scanning electron microscopy (SEM) confirmed biofilm formation on the electrodes’ surface for the three tested anammox bacteria (Fig. S5). The biofilm cell density of MECs inoculated with *Ca*. Brocadia was higher at 0.4 V (Fig. S5E and F) compared to other set potentials, and no biofilm was observed at set potentials ≤ 0 V vs Ag/AgCl (Fig. S5A). These observations correlate very well with the obtained current profiles at different set potentials (Fig. 2A). Cell appendages between cells and the electrode were not observed. Cell appearance was very similar to reported SEM images of anammox cells (*21*).

FISH with anammox-specific probes (Fig. 2F) and metagenomics of DNA extracted from the biofilm on the working electrodes of MECs showed that anammox were the most abundant bacteria in the biofilm community (Fig. S2B and D). Also, no other known electrochemically active bacteria were detected in the metagenomes. Similarly, AOB were not detected which further supports the lack of ATU inhibition on NH_4_^+^ removal and current generation. By differential coverage and sequence composition-based binning (*23*) it was possible to extract high-quality genomes of *Brocadia* and *Scalindua* species from the electrodes (Fig. S2B and 2D). Based on the differences in the genome content, average amino acid identity (AAI) ≤95% compared to reported anammox genomes to date, and evolutionary divergence in phylogenomics analysis (Fig. S6) we propose a tentative name for *Ca*. Brocadia present in our MECs: *Candidatus* Brocadia electricigens (etymology: L. adj. *electricigens*; electricity generator).

## Molecular mechanism of EET-dependent anammox process

To better understand how NH_4_^+^ is converted to N_2_ by anammox bacteria in electrode-dependent anammox process, isotope labelling experiments were carried out. Complete oxidation of NH_4_^+^ to N_2_ was demonstrated by incubating the MECs with ^15^NH_4_^+^ (4 mM) and ^14^NO_2_^−^ (1 mM). Consistent with expected anammox activity, anammox bacteria consumed first the ^14^NO_2_^−^ resulting in the accumulation of ^29^N_2_ in the headspace of the MECs. Interestingly, after depletion of available ^14^NO_2_^−^, a steady increase of ^30^N_2_ was observed with slower activity rates compared to the typical anammox process (Fig 3A, Table S2). These results confirm the GO experiments where ^30^N_2_ was detected when the three anammox species were incubated with ^15^NH_4_^+^ (Fig. 1C). Gas production was not observed in the abiotic control incubations. In the current model of the anammox reaction (Eq. 2) (*4*), NH_4_^+^ is converted to N_2_ with NO_2_^−^ as terminal electron acceptor. This is a process in which first, NO_2_^−^ is reduced to nitric oxide (NO, Eq. 3) and subsequently condensed with ammonia (NH_3_) to produce hydrazine (N_2_H_4_, Eq. 4), which is finally oxidized to N_2_ (Eq. 5). The four low-potential electrons released during N_2_H_4_ oxidation fuel the reduction reactions (Eq. 3 and 4), and are proposed to build up the membrane potential and establish a proton-motive force across the anammoxosome membrane driving the ATP synthesis.

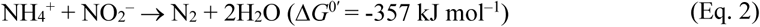

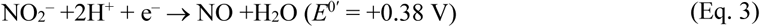

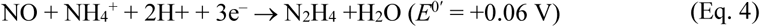

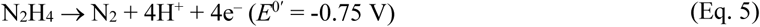

**Figure 3.**
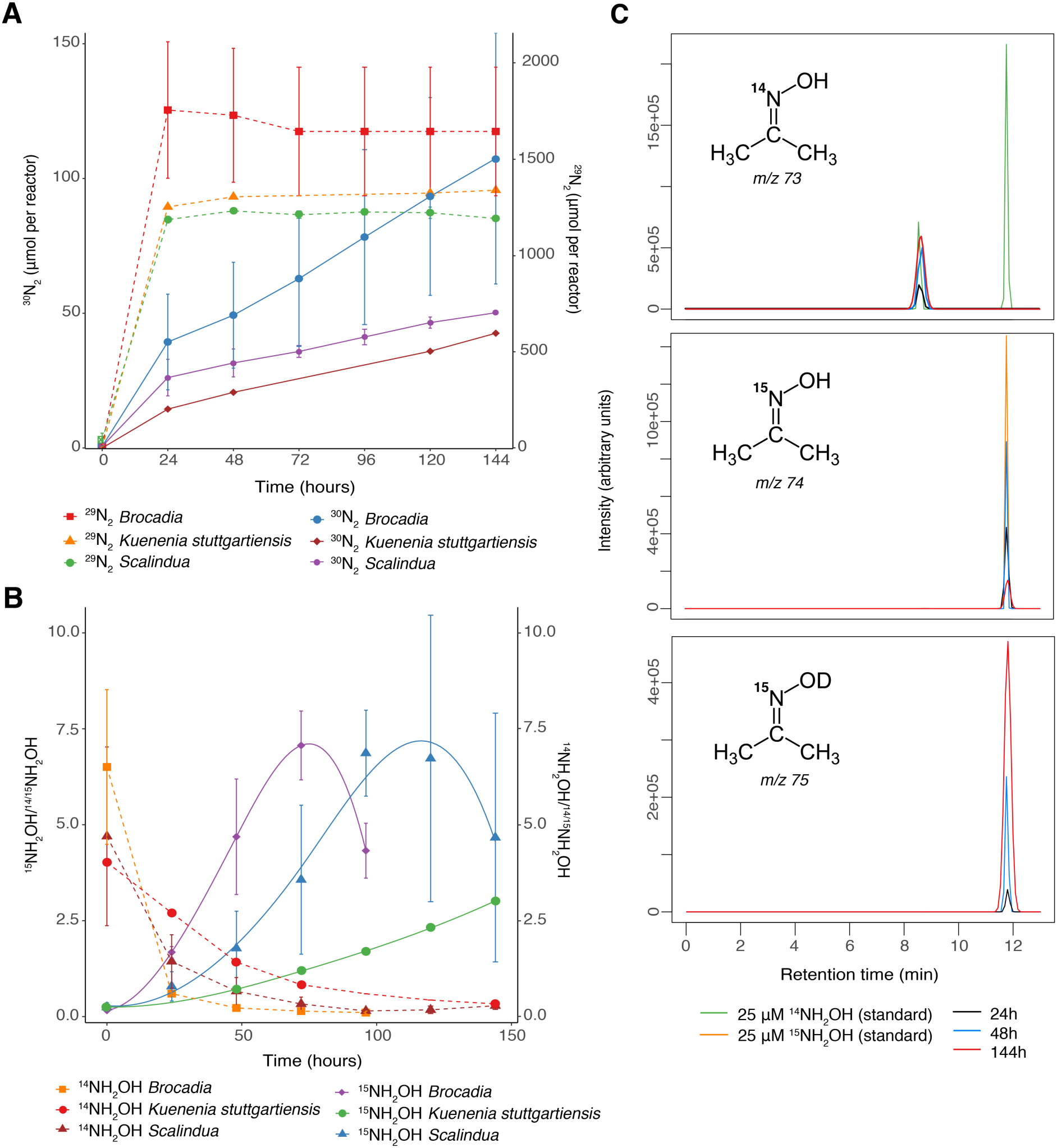
Molecular mechanism of electrode-dependent anaerobic ammonium oxidation by different anammox bacteria. (**A**) Time course of the anaerobic oxidation of ^15^NH_4_^+^ to ^29^N_2_ and ^30^N_2_. The single-chamber MECs with mature biofilm on the working electrodes operated at 0.4 V vs Ag/AgCl were fed with 4 mM ^15^NH_4_^+^ and 1 mM ^14^NO_2_^-^. Under these conditions, anammox bacteria will consume first the preferred electron acceptor (i.e., ^14^NO_2_^-^) and form ^29^N_2_ and then the remaining ^15^NH_4_^+^ will be oxidized to the final product (^30^N_2_) through the electrode-dependent anammox process. NO and N_2_O were not detected throughout the experiment. Results from triplicate MEC reactors are presented as mean ± SD. (**B**) Determination of NH_2_OH as the intermediate of the electrode-dependent anammox process. The MECs with mature biofilm on the working electrodes operated at 0.4 V vs Ag/AgCl were fed with 4 mM ^15^NH_4_^+^ and 2 mM ^14^NH_2_OH. Under these conditions, anammox bacteria would preferentially consume the unlabelled pool of hydroxylamine (i.e., ^14^NH_2_OH), leading to the accumulation of ^15^NH_2_OH due to the oxidation of ^15^NH_4_^+^. Samples were derivatized using acetone, and isotopic ratios were determined by gas chromatography mass spectrometry (GC/MS). Results from triplicate MEC reactors are presented as mean ± SD. (**C**) Ion mass chromatograms of hydroxylamine derivatization with acetone. The MECs with mature biofilm (*Ca.* Brocadia) on the working electrodes operated at 0.4 V vs Ag/AgCl were fed with 4 mM ^15^NH_4_^+^ and 10% D_2_O. The mass to charge (*m/z*) of 73, 74 and 75 corresponds to derivatization products of ^14^NH_2_OH, ^15^NH_2_OH, and ^15^NH_2_OD, respectively with acetone determined by GC/MS. 25 μM of ^14^NH_2_OH and ^15^NH_2_OH were used as standards. The 73 *m/z* (top) at retention time of 8.6 minutes arises from the acetone used for derivatization. The 75 *m/z* (bottom) accumulation over the course of the experiment indicates that the oxygen used in the anaerobic oxidation of ammonium originates from OH^-^ of the water molecule.

In the MEC experiments with *Ca.* Brocadia using electrodes as sole electron acceptor we observed the production of NH_2_OH followed by a transient accumulation of N_2_H_4_ (Fig. S7). No inhibitory effect was observed in incubations with 2-phenyl-4,4,5,5,-tetramethylimidazoline-1-oxyl-3-oxide (PTIO) (Fig. S8), an NO scavenger (*4*). Therefore, we hypothesized that NH_2_OH, and not NO, is an intermediate of the electrode-dependent anammox process. To investigate whether NH_2_OH could be produced directly from NH_4_^+^ in electrode-dependent anammox process, MECs were incubated with ^15^NH_4_^+^ (4 mM) and ^14^NH_2_OH (2 mM). The isotopic composition of the reactors revealed that unlabeled ^14^NH_2_OH was used as a pool substrate and we detected newly synthetized ^15^NH_2_OH from ^15^NH_4_^+^ oxidation (Fig. 3B). It is known that NO and NH_2_OH, the known intermediates in the anammox process, are strong competitive inhibitors of the N_2_H_4_ oxidation activity by the hydrazine dehydrogenase (HDH) (*24*). However, oxidation of N_2_H_4_ (Fig. S7) and detection of ^30^N_2_ (Fig. 3A) in our experiments, suggest that even though there might be some inhibition caused by the NH_2_OH, the HDH is still active. Also, comparative transcriptomics analysis of the electrode’s biofilm revealed that the HDH was one of the most upregulated genes when the electrode was used as the electron acceptor instead of NO_2_^−^ (Supplementary text). Incubations with ^15^NH_4_^+^ (4 mM) in 10% deuterium oxide (D_2_O) showed accumulation of ^15^NH_2_OD, which suggests that in order to oxidize the NH_4_^+^ to NH_2_OH, the different anammox bacteria use OH^−^ ions generated from water (Fig. 3C). Abiotic incubations did not show any production of NH_2_OH or NH_2_OD. Based on these results we propose the following reactions for electrode-dependent anammox process:

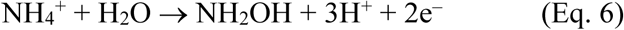

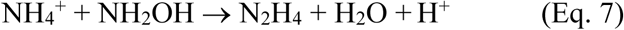

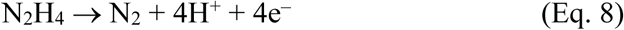

The complete NH_4_^+^ oxidation to N_2_ coupled with reproducible current production can only be explained by electron transfer from the anammoxosome compartment (energetic central of anammox cells and where the NH_4_^+^ is oxidized) to the electrode. In order to compare the pathway of NH_4_^+^ oxidation and electron flow through compartments (anammoxosome) and membranes (cytoplasm and periplasm) in EET-dependent anammox process (electrode poised at 0.4 V vs Ag/AgCl as electron acceptor) versus typical anammox process (i.e., nitrite used as electron acceptor), we conducted a genome-centric comparative transcriptomics analysis (Supplementary text). In the anammoxosome compartment, the genes encoding for ammonium transporter (AmtB), a hydroxylamine oxidoreductase (HAO) and HDH were the most upregulated in response to the electrode as the electron acceptor (Table S8). This observation agrees with the NH_4_^+^ removal and oxidation to N_2_ observed in the MECs and isotope labeling experiments (Fig. 2A, Fig 3A). The genes encoding for NO and NO_2_^−^ reductases (*nir* genes) and their redox couples were significantly downregulated when electrode was used as the electron acceptor (Table S8). This is expected as NO_2_^−^ was not added in the electrode-dependent anammox process. Also, this supports the hypothesis that NO is not an intermediate of the electrode-dependent anammox process and that there was no effect of PTIO when NO_2_^−^ was replaced by the electrode as electron acceptor (Fig. S8). Isotope labelling experiments revealed that NH_2_OH was the intermediate in EET-dependent anammox process and NO was not detected throughout the experiment (Fig. 3B), suggesting that the production of NH_2_OH was not through NO reduction. This was further supported by the observation that the electron transfer module (ETM) and its redox partner whose function is to provide electrons to the hydrazine synthase (HZS) for NO reduction to NH_2_OH were downregulated (Table S6). Interestingly, our analysis revealed that the electrons released from the N_2_H_4_ oxidation (Eq. 8) are transferred to the electrode via an EET pathway that is analog to the ones present in metal-reducing organisms such as *Geobacter* spp. and *Shewanella* spp. (Fig. S10, Supplementary text). Highly expressed cytoplasmic electron carriers such as NADH and ferredoxins can be oxidized at the cytoplasmic membrane by NADH dehydrogenase (NADH-DH) and/or formate dehydrogenase (FDH) to directly reduce the menaquinone pool inside the cytoplasmic membrane (Supplementary materials, Table S3). An upregulated protein similar to CymA (tetraheme c-type cytochrome) in *Shewanella* would then oxidize the reduced menaquinones, delivering electrons to highly upregulated periplasmic cytochromes shuttles and to a porin-cytochrome complex that spans the outer membrane (Fig. S10, Table S3, Supplementary materials). From this complex, electrons could be directly accepted by the insoluble extracellular electron acceptor. Taken together, the results from the comparative transcriptomics analysis suggest an alternative pathway for NH_4_^+^ oxidation coupled to EET when working electrode is used as electron acceptor compared to NO_2_^−^ as electron acceptor.

In conclusion, our study provides the first experimental evidence that phylogenetically and physiologically distant anammox bacteria have EET capability and can couple the oxidation of NH_4_^+^ with transfer of electrons to carbon-based insoluble extracellular electron acceptors. The prevalence of EET-based respiration has been demonstrated using bioelectrochemical systems for both Gram-positive and Gram-negative bacteria (*13, 25*). However, compared to reported EET-capable bacteria, to externalize electrons anammox bacteria have to overcome an additional electron transfer barrier: the anammoxosome compartment. Electrochemically active bacteria are typically found in environments devoid of oxygen or other soluble electron acceptors (*25*). Our results show a novel process of anaerobic ammonium oxidation coupled to EET-based respiration of carbon-based insoluble extracellular electron acceptor by both freshwater and marine anammox bacteria and suggest that this process may also occur in natural anoxic environments where soluble electron acceptors are not available. These results offer a new perspective of a key player involved in the biogeochemical nitrogen cycle. Therefore, a better understanding of EET processes contributes to our understanding of the cycles that occur on our planet (*25*).

## Funding

This work was supported by Center Competitive Funding Program (FCC/1/1971-33-01) to Pascal E. Saikaly from King Abdullah University of Science and Technology (KAUST). Mike S. M. Jetten was supported by ERC AG 232937 and 339880 and SIAM OCW/NWO 024002002.

## Author contributions

D.R.S executed the experiments and analyzed the data. M.A enriched planktonic cells Ca. Brocadia and Scalindua in the MBRs and contributed with the isotope labelling experiments. Bioelectrochemical analysis were done by D.R.S and K.P.K. R.M. and L.V.N designed and executed the scanning electron microscopy analyses. D.R.S performed the metagenomics analysis. D.R.S and M.A. did the phylogenomics analysis. D.R.S and M.S.M.J. designed the Isotopic batch experiments. D.R.S and J.R. did the comparative transcriptomics analysis and developed the molecular model. D.R.S., M.A., K.P.K., M.S.M.J and P.E.S. planned the research. D.R.S wrote the paper with critical feedback from P.E.S., M.S.M.J., L.V.N, J.A.G, M.A., K.P.K., J.R., and R.M.

## Competing interests

Authors declare no competing interests.

## Data and materials availability

The genome binning and the comparative transcriptomics analysis are entirely reproducible using the R files available on https://github.com/DarioRShaw/Electro-anammox. Also, complete Datasets generated in the differential expression analysis are available in the online version of the paper. Raw sequencing reads of Illumina HiSeq of metagenomics and metatranscriptomics data associated with this project can be found at the NCBI under BioProject PRJNA517785. Annotated GenBank files for the anammox genomes extracted in this study can be found under the accession numbers SHMS00000000 and SHMT00000000.

## Supplementary Materials

### Materials and Methods

#### Enrichment and cultivation of anammox bacteria

Biomass from upflow column reactors (XK 50/60 Column, GE Healthcare, UK) with *Ca*. Brocadia and *Ca*. Scalindua were harvested and used as inoculum. *Ca*. Brocadia and *Ca*. Scalindua planktonic cells were enriched in two bioreactors (BioFlo®115, New Brunswick, USA) equipped with a microfiltration (average pore size 0.1 µm) hollow fiber membrane module (zena-membrane, Czech Republic) (Fig. S1A). Operating conditions of the membrane bioreactors (MBRs) were described previously (*15*). The MBRs were operated at pH 7.5–8.0 and 35±1°C for *Brocadia* and room temperature (20–25 C) for *Scalindua*. The culture liquid in the MBRs was continuously mixed with a metal propeller at a stirring speed of 150 rpm and purged with 95% Ar – 5% CO_2_ at a flow rate of 10 mL min^−1^ to maintain anaerobic conditions. Inorganic synthetic medium was fed continuously to the reactors at a rate of ∼5 L d^−1^ and hydraulic retention time was maintained at one day. The synthetic medium was prepared by adding the following constituents; NH_4_^+^ (2.5-10) mM, NO_2_^−^ (2.5-12) mM, CaCl_2_ 100 mg L^−1^, MgSO_4_ 300 mg L^−1^, KH_2_PO_4_ 30 mg L^−1^, KHCO_3_ 500 mg L^−1^ and trace element solutions (*25*). In the case of *Ca*. Scalindua culture, the synthetic medium was prepared using non-sterilized Red Sea water. Samples for microbial community characterization were taken from the MBRs for fluorescence *in situ* hybridization (FISH) and metagenomics analysis (See FISH and DNA extraction, metagenome library preparation, sequencing and sequence processing and analysis sections below). A previously enriched *K. stuttgartiensis* culture was also used for the experiments (*4*).

#### Incubation of anammox bacteria in serum vials with NH_4_^+^ and graphene oxide as the insoluble extracellular electron acceptor

To test whether anammox bacteria have extracellular electron transfer (EET) capability, the three enriched anammox cultures were incubated in serum vials for 216 hours with ^15^NH_4_^+^ and graphene oxide (GO) as a proxy for insoluble extracellular electron acceptor. Standard anaerobic techniques were employed in the batch incubation experiments. All the procedures were performed in the anaerobic chamber (Coy Laboratory Products; Grass Lake Charter Township, MI, USA). Anoxic buffers and solutions were prepared by repeatedly vacuuming and purging helium gas (>99.99%) before experiments. Biomass from the MBRs was centrifuged, washed twice and suspended in inorganic medium containing 2 mM 4-(2-hydroxyethyl)-1-piperazineethanesulfonic acid (HEPES, pH 7.8) prior to inoculation into the vials. The same composition of the inorganic medium used in the MBRs was supplied to the vials. Cell suspension was dispensed into 100 mL glass serum vials, which were sealed with butyl rubber stoppers and aluminum caps. Biomass concentration in the vials ranged from 0.1–0.9 mg–protein mL^−1^. The headspace of the serum vials was replaced by repeatedly vacuuming and purging with pure (>99.99%) helium gas. Positive pressure (50–75 kPa) was added to the headspace to prevent unintentional contamination with ambient air during the incubation and gas sampling. Prior the addition of ^15^NH_4_^+^, the vials were pre-incubated overnight at room temperature (∼25°C) to remove any trace amounts of substrates and oxygen. The activity test was initiated by adding 4 mM of ^15^NH_4_Cl (Cambridge Isotope Laboratories) and GO to a final concentration of 200 mg L^−1^ using a gas–tight syringe (VICI; Baton Rouge, LA, USA). No NO_2_^−^ or NO_3_^−^ were added to the incubations. The vials were incubated in triplicates at 30°C for *Ca*. Brocadia and *K. stuttgartiensis* cultures and at room temperature (∼25°C) for vials with *Ca*. Scalindua. Vials without biomass were also prepared as abiotic controls. The concentrations of ^28^N_2_, ^29^N_2_ and ^30^N_2_ gas were determined by gas chromatography mass spectrometry (GC/MS) analysis (*27*). Fifty microliter of headspace gas was collected using a gas-tight syringe (VICI; Baton Rouge, LA, USA) and immediately injected into a GC (Agilent 7890A system equipped with a CP-7348 PoraBond Q column) combined with 5975C quadrupole inert MS (Agilent Technologies; Santa Clara, CA, USA), and mass to charge (m/z) = 28, 29 and 30 was monitored. Standard calibration curve of N_2_ gas was prepared with ^30^N_2_ standard gas (>98% purity) (Cambridge Isotope Laboratories; Tewksbury, MA, USA). At the end of the batch incubations, DNA was extracted and sequenced for metagenomics analysis (See DNA extraction, metagenome library preparation, sequencing and sequence processing and analysis section below). To confirm the reduction of the GO, the samples were centrifuged and subjected to dehydration process with absolute ethanol. Samples were maintained in a desiccator until Raman spectroscopy analysis. Raman spectroscopy (StellarNet Inc) was performed with the following settings: Laser 473 nm, acquisition time 20 seconds, accumulation 5 and objective 50X.

#### Bioelectrochemical analyses

To evaluate if anammox bacteria (*Ca*. Brocadia and *Ca*. Scalindua) are electrochemically active, single-chamber multiple working electrode glass reactors with 500 mL working volume were operated in microbial electrolysis cell (MEC) mode. The working electrodes (anodes) were graphite rods of 8 cm length (7.5 cm inside the reactor) and 0.5 cm in diameter. Platinum mesh was used as counter electrode (cathode) and Ag/AgCl as reference electrode (Bioanalytical Systems, Inc.). A schematic representation of the multiple working electrode microbial electrolysis cell (MEC) is presented in Fig. S1H. The multiple working electrodes were operated at a set potential of –0.3, –0.2, –0.1, 0, 0.1, 0.2, 0.3 and 0.4 V vs Ag/AgCl. Amperometric current was monitored continuously using a VMP3 potentiostat (BioLogic Science Instruments, USA), with measurements every 60 s and analyzed using EC-lab V 10.02 software. To evaluate if *K. stuttgartiensis* is electrochemically active, experiments were conducted in single-chamber MECs (300 mL working volume) with carbon cloth working electrode (0.4 V vs Ag/AgCl). The reactors and working and counter electrodes were sterilized by autoclaving prior to the start of the experiments. The reference electrodes were sterilized by soaking in 3 M NaCl overnight and rinsing with sterile medium. After the reactors were assembled, epoxy glue was used to seal every opening in the reactor to avoid leakage. Gas bags (0.1 L Cali −5-Bond. Calibrate, Inc.) were connected to the MECs to collect any gas generated. The gas composition in the gas bags was analyzed using a gas chromatograph (SRI 8610C gas chromatograph, SRI Instruments).

The inorganic medium composition in the MECs was the same as the one supplied in the MBRs (See Enrichment and cultivation of anammox bacteria section above), with variations in the NH_4_^+^ and/or NO_2_^−^ concentration. After preparation, the inorganic medium was boiled, sparged with N_2_:CO_2_ (80:20) gas mix for 30 min to remove any dissolved oxygen and finally autoclaved. The autoclaved medium was cooled down to room temperature inside the anaerobic chamber (Coy Laboratory, USA). Prior to the experiments, KHCO_3_ was weighed in the anaerobic chamber and dissolved in the medium. The reactors were operated in fed batch mode at 30°C for *Ca*. Brocadia and *K. stuttgartiensis* cultures and at room temperature (∼25°C) for *Ca*. Scalindua. The medium in the MECs was gently mixed with a magnetic stirrer throughout the course of the experiments. The pH of the MECs was not controlled but was at all times between 7.0–7.5. To exclude the effect of abiotic (i.e., non-Faradaic) current, initial operation of the reactors was done without any biomass addition. After biomass inoculation, the MECs were operated with set potentials and optimal conditions for the anammox reaction (i.e., addition of NH_4_^+^ and NO_2_^−^). Afterwards, NO_2_^−^ was gradually decreased to 0 mM leaving the working electrodes as the sole electron acceptor. To confirm that the electrode-dependent anaerobic oxidation of NH_4_^+^ was catalyzed by anammox bacteria, additional control experiments were conducted in chronological order including addition of allylthiourea (ATU), operation in open circuit voltage mode (i.e., anodes were not connected to the potentiostat; electrode is not used as electron acceptor), addition of nitrite, operation without addition of NH_4_^+^ and then with addition of NH_4_^+^, and autoclaving. ATU was added to a final concentration of 100 μM to evaluate the contribution of nitrifiers to the process (*19*). Biomass from a nitrifying reactor was incubated in triplicate vials with 100 μM of ATU and was used as a positive control for the inhibitory effect of ATU. Throughout the reactor operation, the concentrations of NH_4_^+^, NO_2_^−^, and NO_3_^−^ were determined as described below (See Analytical methods section). All experiments were done in triplicate MECs, unless mentioned otherwise.

Cyclic voltammetry (CV) at a scan rate of 1mV s^−1^ was performed for the anodic biofilms at different time intervals following initial inoculation to determine their redox behavior. Scans ranged from −0.8 to 0.4 V vs Ag/AgCl. Current was normalized to the geometric anode surface area. To determine the presence of extracellular secreted redox mediators by anodic communities, CVs were performed with cell-free filtrates (filtered using a 0.2 mm pore diameter filter) collected from the reactors and placed in separate sterile electrochemical cells. Also, experiments were conducted to evaluate the effect of adding riboflavin, which is a common soluble mediator involved in EET in gram-positive and gram-negative bacteria (*13, 20*). Riboflavin was added to the mature anammox biofilm to a final concentration of 250 nM (*20*).

To test if cathodic reaction (i.e., hydrogen evolution reaction) has an effect on electrode-dependent anaerobic ammonium oxidation, experiments were also conducted in double-chamber MECs (Fig. S1K) with a single carbon cloth working electrode (0.4 V vs Ag/AgCl). The anode and cathode chambers in double-chamber MECs were separated by a proton-exchange Nafion membrane. Also, to exclude the effect of heterotrophic activity on current generation, 500 mg L^−1^ of penicillin G (Sigma-Aldrich, St. Louis, MO) was added in the last batch cycle to inhibit heterotrophs (*21, 22*).

To determine the role of NO in the electrode-dependent anammox metabolism, single-chamber MECs were incubated with 4 mM NH_4_^+^ and 100 M of 2-phenyl-4,4,5,5,-tetramethylimidazoline-1-oxyl-3-oxide (PTIO), an NO scavenger. MECs with 4 mM NH_4_^+^ and without PTIO addition were run in parallel as negative control. PTIO inhibits *K. stuttgartiensis* activity when NO is an intermediate of the anammox reaction (*4*), therefore vials with *K. stuttgartiensis* were used as positive control of the effect of PTIO. Liquid samples were taken every day and filtered using a 0.2 mm filter and subjected to determination of NH_4_^+^ concentration as described below (See Analytical methods section).

For isotopic and comparative transcriptomics analysis experiments, single-chamber MECs (Adams & Chittenden Scientific Glass, USA) with a single carbon cloth working electrode (0.4 V vs Ag/AgCl) and 300 mL working volume were used (Fig. S1J).

#### 15N tracer batch experiments in MECs

To elucidate the molecular mechanism of electrode-dependent anaerobic ammonium oxidation by different anammox bacteria, isotopic labelling experiments were conducted in single-chamber MECs operated at set potential of 0.4 V vs Ag/AgCl. All batch incubation experiments were performed in triplicate MECs. MEC incubations without biomass for the ^15^N tracer batch experiments were also prepared to exclude any possibility of an abiotic reaction. Standard anaerobic techniques were employed in the batch incubation experiments. All the procedures were performed in the anaerobic chamber (Coy Laboratory Products; Grass Lake Charter Township, MI, USA). Anoxic buffers and solutions were prepared by repeatedly vacuuming and purging helium gas (>99.99%) before the experiments. Purity of ^15^N-labelled compounds was greater than 99%. The headspace of the MECs was replaced by repeatedly vacuuming and purging with pure (>99.99%) helium gas. Positive pressure (50–75 kPa) was added to the headspace to prevent unintentional contamination with ambient air during the incubation and gas sampling. Oxidation of NH_4_^+^ to N_2_ was demonstrated by incubating the MECs with ^15^NH_4_Cl (Cambridge Isotope Laboratories, 4 mM) and ^14^NO_2_^−^ (1 mM). The MECs were incubated for 144 hours at 30°C for *Ca*. Brocadia and *K. stuttgartiensis* cultures, and at room temperature (∼25°C) for *Ca*. Scalindua. The concentrations of ^28^N_2_, ^29^N_2_, ^30^N_2_, ^14^NO, ^15^NO, ^28^N_2_O, ^29^N_2_O and ^30^N_2_O gas were determined by GC/MS (*27*). Fifty microliter of headspace gas was collected using a gas-tight syringe (VICI; Baton Rouge, LA, USA) and immediately injected into a GC (Agilent 7890A system equipped with a CP-7348 PoraBond Q column) combined with 5975C quadrupole inert MS (Agilent Technologies; Santa Clara, CA, USA). Standard calibration curve of N_2_ gas was prepared with ^30^N_2_ standard gas (>98% purity) (Cambridge Isotope Laboratories; Tewksbury, MA, USA).

To investigate whether hydroxylamine (NH_2_OH) could be produced directly from NH_4_^+^ in electrode-dependent anaerobic ammonium oxidation by anammox bacteria, single-chamber MECs were incubated with ^15^NH_4_Cl (4 mM, Cambridge Isotope Laboratories) and an unlabeled pool of ^14^NH_2_OH (2 mM) for 144 hours. Liquid samples were taken every day and filtered using a 0.2 mm filter and subjected to determination of ^15^NH_2_OH and ^14^NH_2_OH. NH_2_OH was determined by GC/MS analysis after derivatization using acetone (*27*). Briefly, 100 µl of liquid sample was mixed with 4 µl of acetone, and 2 µl of the derivatized sample was injected to a GC (Agilent 7890A system equipped with a CP-7348 PoraBond Q column) combined with 5975C quadrupole inert MS (Agilent Technologies; Santa Clara, CA, USA) in splitless mode. NH_2_OH was derivatized to acetoxime (C_3_H_7_NO), and the molecular ion peaks were detected at mass to charge (m/z) = 73 and 74 for ^14^NH_2_OH and ^15^NH_2_OH, respectively. 25 M of ^14^NH_2_OH and ^15^NH_2_OH were used as standards. To determine the source of the oxygen used in the electrode-dependent NH_4_^+^ oxidation to NH_2_OH, MECs were incubated with ^15^NH_4_Cl (4 mM, Cambridge Isotope Laboratories) in presence of 10% D_2_O for 144 hours. Stable isotopes of NH_2_OH were determined by GC/MS analysis after derivatization using acetone as described above.

#### Activity and electron balance calculations

Activities of specific anammox (^29^N_2_) with nitrite as the preferred electron acceptor and electrode-dependent anammox (^30^N_2_) with working electrode (0.4 V vs Ag/AgCl) as sole electron acceptor were calculated based on the changes in gas concentrations in single-chamber MEC batch incubations. The activity was normalized against protein content of the biofilm on the electrodes. Protein content was measured as described below (See Analytical methods section).

The moles of electrons recovered as current per mole of NH_4_^+^ oxidized were calculated using:

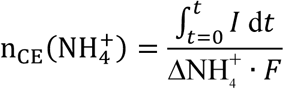

where *I* is the current (A) obtained from the chronoamperometry, d*t* (s) is the time interval over which data was collected, NH_4_^+^ is the moles of NH_4_^+^ consumed during the experiment, and *F* = 96485 C/mol e^−^ is Faraday’s constant. Coulombic efficiency (CE) was calculated using:

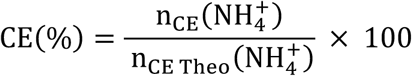

where n_CE Theo_(NH_4_^+^) is the theoretical number of moles of electrons (in our case it is 3 moles of electrons) recovered as current per mole of NH_4_^+^ oxidized.

#### Analytical methods

All samples were filtered through a 0.2 µm pore-size syringe filters (Pall corporation) prior to chemical analysis. NH_4_^+^ concentration was determined photometrically using the indophenol method (*28*) (lower detection limit = 5 μM). Absorbance at a wavelength of 600 nm was determined using multi-label plate readers (SpectraMax Plus 384; Molecular Devices, CA, USA). NO_2_^−^ concentration was determined by the naphthylethylenediamine method (*28*) (lower detection limit = 5 μM). Samples were mixed with 4.9 mM naphthylethylenediamine solution, and the absorbance was measured at a wavelength of 540 nm. NO_3_^−^ concentration was measured by HACH kits (HACH, CO, USA; lower detection limit = 0.01 mg l^-1^ NO_3_^-^-N). User’s guide was followed for these kits and concentrations were measured by spectrophotometer (D5000, HACH, CO, USA). Concentrations of NH_2_OH and hydrazine (N_2_H_4_) were determined colorimetrically as previously described (*29*). For NH_2_OH, liquid samples were mixed with 8-quinolinol solution (0.48% (w/v) trichloroacetic acid, 0.2% (w/v) 8-hydroxyquinoline and 0.2 M Na_2_CO_3_) and heated at 100°C for 1 min. After cooling down for 15 min, absorbance was measured at 705 nm (*30*). N_2_H_4_ was derivatized with 2% (w/v) p-dimethylaminobenzaldehyde and absorbance at 460 nm was measured (*31*). The concentration of biomass on the working electrodes was determined as protein concentration using DC Protein Assay Kit (Bio-Rad, Tokyo, Japan) according to manufacturer’s instructions. Bovine serum albumin was used as the protein standard.

#### Fluorescence *in situ* hybridization

The microbial community in the MBRs and the spatial distribution of anammox cells on the surface of the graphite rod electrodes was examined by FISH after 30 days of reactor operation. The graphite rod electrodes were cut in the anaerobic chamber with sterilized tube cutter (Chemglass Life Sciences, US). The electrode samples were fixed with 4% (v/v) paraformaldehyde (PFA), followed by 10 nm cryosectioning at −30°C (Leica CM3050 S Cryostat). FISH with rRNA-targeted oligonucleotide probes was performed as described elsewhere (*32*) using the EUB338 probe mix composed of equimolar EUB338 I, EUB338 II and EUB 338 III (*33, 34*) for the detection of bacteria and probes AMX820 or SCA1309 for anammox (*35, 36*). Cells were counterstained with 1 μg ml^−1^ DAPI (4′,6-diamidino-2-phenylindole) solution. Fluorescence micrographs were recorded by using a Leica SP7 confocal laser scanning microscope. To determine the relative abundance of anammox bacteria by quantitative FISH, 20 confocal images of FISH probe signals were taken at random locations in each well and analyzed by using the digital image analysis DAIME software as described elsewhere (*37*).

#### Scanning Electron Microscopy

The graphite rod electrodes were cut in the anaerobic chamber with sterilized tube cutter (Chemglass Life Sciences, US). The electrode samples were soaked in 2% glutaraldehyde solution containing phosphate buffer (50 mM, pH 7.0) and stored at 4°C. Sample processing and scanning electron microscopy (SEM) was performed as described elsewhere (*38*). Samples from the carbon-cloth electrodes were punched out using a 4.8 mm Ø biopsy punch and placed into a 200 μm cavity of a type A platelet (6 mm diameter; 0.1-0.2 mm depth, Leica Microsystems) and closed with the flat side of a type B platelet (6 mm diameter, 300 μm depth). Platelet sandwiches were cryo-fixed by high-pressure freezing (Leica HPM 100; Leica Microsystems, Vienna, Austria) and stored in liquid nitrogen until use. For Hexamethyldisalizane (HMDS) embedding, frozen samples were freeze-substituted in anhydrous methanol containing 2% osmium tetroxide, 0.2% uranyl acetate and 1% H_2_O (*39*). The substitution followed several intervals: cells were kept at −90°C for 47 hours; brought to −60°C at 2°C per hour and kept at −60°C for 8 hours; brought to −30°C at 2°C per hour and kept at −30°C for 8 hours in a freeze-substitution unit (AFS; Leica Microsystems, Vienna, Austria). To remove fixatives the samples were washed four times for 30 min in the AFS device at −30°C with anhydrous methanol and subsequently infiltrated with HMDS by incubating two times for 15 minutes with 50% HMDS in anhydrous methanol followed by two times 15 minutes 100% HMDS. After blotting and air-drying the electrode samples were mounted on specimen stubs using conductive carbon tape and sputter-coated with gold-palladium before imaging in a JEOL JSM-6335F SEM, operating at 3kV.

#### DNA extraction, metagenome library preparation, sequencing and sequence processing and analysis

Biomass from the vials of the GO experiment was harvested by centrifugation (4000g, 4°C) at the end of the batch incubations. Biofilm samples from the electrodes were collected after 30 days of reactor operation with working electrode as the sole electron acceptor. The biomass pellet and the electrode samples were suspended in Sodium Phosphate Buffer in the Lysing Matrix E 2 mL tubes (MP Biomedicals, Tokyo, Japan). After 2 minutes of physical disruption by bead beating (Mini-beadbeater^TM^, Biospec products), the DNA was extracted using the Fast DNA spin kit for soil (MP Biomedicals, Tokyo, Japan) according to the manufacturer’s instructions. The DNA was quantified using Qubit (Thermo Fisher Scientific, USA) and fragmented to approximately 550 bp using a Covaris M220 with microTUBE AFA Fiber screw tubes and the settings: duty factor 20%, peak/displayed power 50W, cycles/burst 200, duration 45s and temperature 20°C. The fragmented DNA was used for metagenome preparation using the NEB Next Ultra II DNA library preparation kit. The DNA library was paired-end sequenced (2 x 301bp) on a Hiseq 2500 system (Illumina, USA).

Raw reads obtained in the FASTQ format were processed for quality filtering using Cutadapt package v. 1.10 (*40*) with a minimum phred score of 20 and a minimum length of 150 bp. The trimmed reads were assembled using SPAdes v. 3.7.1 (*41*). The reads were mapped back to the assembly using minimap2 (*42*) (v. 2.5) to generate coverage files for metagenomic binning. These files were converted to the sequence alignment/map (SAM) format using samtools (*43*). Open reading frames (ORFs) were predicted in the assembled scaffolds using Prodigal (*44*). A set of 117 hidden Markov models (HMMs) of essential single-copy genes were searched against the ORFs using HMMER3 (http://hmmer.janelia.org/) with default settings, with the exception that option (-cut_tc) was used (*45*). Identified proteins were taxonomically classified using BLASTP against the RefSeq protein database with a maximum e-value cut-off of 10^−5^. MEGAN was used to extract class-level taxonomic assignments from the BLAST output (*46*). The script network.pl (http://madsalbertsen.github.io/mmgenome/) was used to obtain paired-end read connections between scaffolds. 16S rRNA genes were identified using BLAST (*47*) (v. 2.2.28+, and the 16S rRNA fragments were classified using SINA (*48*) (v. 1.2.11) with default settings except min identity adjusted to 0.80. Additional supporting data for binning was generated according to the description in the mmgenome package (*49*) (v. 0.7.1.). Genome binning was carried out in R (*50*) (v. 3.3.4) using the R-studio environment. Individual genome bins were extracted using the multimetagenome principles (*23*) implemented in the mmgenome R package (*50*) (v. 0.7.1). Completeness and contamination of bins were assessed using coverage plots through the mmgenome R package and by the use of CheckM (*51*) based on occurrence of a set of single-copy marker genes (*52*). Genome bins were refined manually as described in the mmgenome package and the final bins were annotated using PROKKA (*53*) (v. 1.12-beta).

#### Phylogenomics analysis

Extracted bins and reported anammox genomes were used for phylogenetic analysis. Reported anammox genomes were downloaded from the NCBI GenBank. Hidden Markov model profiles for 139 single-copy core genes (*52*) were concatenated using anvi’o platform (*54*). Phylogenetic trees with estimated branch support values were constructed from these concatenated alignments using MEGA7 (*55*) with Neighbor Joining, Maximum-likelihood and UPGMA methods.

#### Comparative transcriptomics analysis

Comparative transcriptomic analysis was conducted to compare the metabolic pathway of NH_4_^+^ oxidation and electron flow when working electrode is used as electron acceptor versus NO_2_^−^ as electron acceptor. Samples for comparative transcriptomic analysis were taken from mature electrode’s biofilm of duplicate single-chamber MECs with NO_2_^−^ as the sole electron acceptor and after switching to set potential growth (0.4 V vs Ag/AgCl, electrode as electron acceptor). Biofilm samples were collected from carbon cloth electrodes with sterilized scissors in the anaerobic chamber. Samples were stored in RNAlater™ Stabilization Solution (Invitrogen™) until further processing. Total RNA was extracted from the samples using PowerBiofilm RNA Isolation kit (QiAGEN) according to manufacturer’s instructions. The RNA concentration of all samples was measured in duplicate using the Qubit BR RNA assay. The RNA quality and integrity were confirmed for selected samples using TapeStation with RNA ScreenTape (Agilent Technologies). The samples were depleted of rRNA using the Ribo-zero Magnetic kit (Illumina Inc.) according to manufacturer’s instructions. Any potential residual DNA was removed using the DNase MAX kit (MoBio Laboratories Inc.) according to the manufacturer’s instructions. After rRNA depletion and DNase treatment the samples were cleaned and concentrated using the RNeasy MinElute Cleanup kit (QIAGEN) and successful rRNA removal was confirmed using TapeStation HS RNA Screentapes (Agilent Technologies). The samples were prepared for sequencing using the TruSeq Stranded Total RNA kit (Illimina Inc.) according to the manufacturer’s instructions. Library concentrations were measured using Qubit HS DNA assay and library size was estimated using TapeStation D1000 ScreenTapes (Agilent Technologies). The samples were pooled in equimolar concentrations and sequenced on an Illumina HiSeq2500 using a 1×50 bp Rapid Run (Illumina Inc).

Raw sequence reads in fastq format were trimmed using USEARCH (*56*) v10.0.2132, - fastq_filter with the settings -fastq_minlen 45 -fastq_truncqual 20. The trimmed transcriptome reads were also depleted of rRNA using BBDuk (*57*) with the SILVA database as reference database (*58*). The reads were mapped to the predicted protein coding genes generated from Prokka (*53*) v1.12 using minimap2 (*42*) v2.8-r672, both for the total metagenome and each extracted genome bin. Reads with a sequence identity below 0.98 were discarded from the analysis. The count table was imported to R (*50*), processed and normalized using the DESeq2 workflow (*59*) and then visualized using ggplot2. Analyses of overall sample similarity were done using normalized counts (log transformed), through vegan (*60*) and DESeq2 (*59*) packages. Differentially expressed genes were evaluated for the presence of N-terminal signal sequences, transmembrane spanning helices (TMH) and subcellular localization using SignalP 5.0 (*62*), TMHMM 2.0 software and PSORTb 3.0.2 (*62*) respectively. Differentially expressed genes that appeared annotated as ‘hypothetical’ were reconsidered for a putative function employing BLAST searches (i.e., BLASTP, CD-search, SmartBLAST), MOTIF search, COG and PFAM databases, as well as by applying the HHpred homology detection and structure prediction program (MPI Bioinformatics Toolkit).

#### Statistics and reproducibility

The number of replicates is detailed in the subsections for each specific experiment and was mostly determined by the amount of biomass available for the different cultures. In all experiments, three biological replicates were used, unless mentioned otherwise. No statistical methods were used to predetermine the sample size. The experiments were not randomized, and the investigators were not blinded to allocation during experiments and outcome assessment. Statistical analyses were carried out in R (*50*) v. 3.3.4 using the R-studio environment.

### Supplementary Text

#### Putative EET-dependent anammox pathway

We provided evidence that phylogenetically distant anammox bacteria can perform EET and are electrochemically active, and we elucidated the molecular mechanism of NH_4_^+^ oxidation, which by itself are significant findings that changes our perception of a key player in the global nitrogen cycle. Next, we conducted comparative transcriptomic analysis to compare the possible pathways involved in the EET-dependent anammox process (electrode poised at 0.4 V vs Ag/AgCl as electron acceptor) versus typical anammox process (i.e., NO_2_^−^ as electron acceptor). Currently, pure cultures of anammox bacteria are unavailable to conduct mutant studies to address the genetic basis of EET-dependent anammox process (*13*). Also, the slow growth rates of anammox bacteria and the fact that they do not rapidly degrade the majority of their proteins, make the changes to specific conditions (i.e., changes in the electron acceptor) not immediately reflected at the protein level (*5*). Therefore, to detect immediate changes in response to a stimulus, short-term gene expression responses would be more appropriate for this aim. In our study, the potential metabolic pathways involved in EET-dependent anammox process, were studied using a genome-centric stimulus-induced transcriptomics approach that has been successfully applied before to identify metabolic networks within complex EET-active microbial communities (*63*). RNA samples were extracted from mature electrode biofilm of two independent single-chamber *Ca*. Brocadia MECs operated first with NO_2_^−^ as the sole electron acceptor and after switching to set potential growth (0.4 V vs Ag/AgCl), and were subjected to a comparative transcriptomics analysis. Similar experiments were conducted with *Ca*. Scalindua and *K. stuttgartiensis*, but we did not get sufficient mRNA from the biofilm samples, and hence only the data for *Ca*. Brocadia are presented here. High similarity was observed between the biological replicates and differentially expressed genes across the experimental setups (Fig. S9). Based on the known cell biology (*64*), biochemistry and anammox metabolism (*4, 65, 66*), and the expression profiles of the known anammox pathways obtained with the differential expression analysis done in this study (Table S3 and 4), we propose a putative molecular model to describe how electrons flow from the anammoxosome to the electrode in the EET-dependent anammox process (Fig. S10). The most differentially expressed genes in response to the change to electrode as electron acceptor were mainly associated with energy conservation and nitrogen metabolism (Table S6 and 8). The metabolic challenge that must be solved for EET process in anammox cells is to transfer the electrons through the separate compartments and membranes (anammoxosome, cytoplasm and periplasm). The observed NH_4_^+^ oxidation and reproducible current generation can only be explained by electrons being transported from the anammoxosome (energetic central of the cell and where the NH_4_^+^ is oxidized) to the electrode. In the anammoxosome, the genes encoding for ammonium transporters (AmtB), a hydroxylamine oxidoreductase (HAO) and hydrazine dehydrogenase (HDH) were the most upregulated (Fig. S10, Table S8). This result is consistent with the NH_4_^+^ uptake, oxidation and final conversion to N_2_ observed in the MECs and isotope labeling experiments (Fig. 2A, Fig 3A). The requirement of more moles of NH_4_^+^ when anammox growth is based on EET compared to NO_2_^−^ as electron acceptor (Eq. 1), increases the demand of NH_4_^+^ import into the cell, which can explain the upregulation of the ammonium transporters. In contrast, the genes encoding for NO and NO_2_^−^ reductases (*nir* genes) and their redox couples were significantly downregulated (Fig. S10, Table S8). This agrees, with the fact that NO_2_^−^ and NO_3_^−^ were below the detection limit in the MECs (Fig. 2A, Fig. S3A and B) and there was no effect of PTIO when NO_2_^−^ was replaced by electrode as electron acceptor (Fig. S8). Also, this supports the hypothesis that NO is not an intermediate of the electrode-dependent anammox process. The most downregulated HAO in the electrode-dependent anammox process (EX330_09385, Table S8) is an ortholog of the proposed nitrite reductase in *K. stuttgartiensis* kustc0458 (*66*). Currently, nitrite reductase(s) responsible for NO_2_^−^ reduction in *Brocadia* species are unidentified (*29*). Therefore, it would be of interest to further investigate the function of the downregulated HAO found in this study as possible candidate for *nir* in *Brocadia*. On the other hand, the *nxr* genes encoding for the soluble nitrite:nitrate oxidoreductase maintained similar levels of expression under both conditions (Table S9). However, cytochromes of the *nxr* gene cluster and the hypothetical membrane-bound NXR were found downregulated under set-potential (Table S8). Even though ammonia is difficult to activate under anaerobic conditions (*67*), previous studies have reported anaerobic NH_4_^+^ oxidation in bioelectrochemical systems dominated by nitrifiers (*68–73*), but the molecular mechanism was not elucidated. Also, an alternative process to anammox called Feammox has been reported recently, where NH_4_^+^ oxidation is coupled with Fe(III) reduction by the Actinobacteria *Acidimicrobiaceae* sp. A6 (*74, 75*). It should be noted that *Acidimicrobiaceae* sp. A6 is not recognized as a key player in the nitrogen cycle. When pure culture of *Acidimicrobiaceae* sp. A6 was tested in MECs with electrode as electron acceptor, there was no colonization and biofilm formation over the course of the experiment. The majority of *Acidimicrobiaceae* cells were present in suspension in the MECs, which explains the low Coulombic Efficiency of the process (∼16.4%) and the need for the soluble electron shuttle 9,10-anthraquinone-2,6-disulfonic acid (AQDS). In the absence of AQDS, no change in NH4^+^ concentration was detected. Future experiments are needed to differentiate Fe(III) reduction for nutritional acquisition from respiration through EET, and to address the genetic basis of the Feammox process and elucidate the molecular mechanism of NH_4_^+^ oxidation. Our isotope labelling experiments revealed that NH_2_OH is a key intermediate in the oxidation of NH_4_^+^ in electrode-dependent anammox process (Fig. 3B), suggesting that the internalized NH_4_^+^ is oxidized to NH_2_OH. More than 10 paralogs of HAO-like proteins in anammox are the most likely candidate enzymes catalyzing anaerobic NH_4_^+^ oxidation. The only upregulated HAO-like protein (EX330_11045) (Fig. S10, Table S8), whose function is still uncharacterized, lacks the tyrosine residue needed for crosslinking of catalytic heme 4, thereby favoring reductive reactions (*29*). This HAO is an ortholog of *K. stuttgartiensis* kustd2021 which under normal anammox conditions has low expression levels (*76*). However, it is worth mentioning that under set potential the whole gene cluster EX330_11030-11050 was significantly upregulated (Table S10). Thus, further investigation should focus on determining the role of this cluster in electrode-dependent anammox process. The produced NH_2_OH is then condensed with NH_3_ to produce N_2_H_4_ by the hydrazine synthase (HZS) (*77*) (Fig. S10). Recent crystallography study of *Ca*. *K. stuttgartiensis* HZS, suggested that N_2_H_4_ synthesis is a two-step reaction: NO reduction to NH_2_OH and subsequent condensation of NH_2_OH and NH_3_ (*77*). Our isotope labelling experiments showed that NH_2_OH is an intermediate in the electrode-dependent anammox process, and thus there is no need for the reduction of NO to NH_2_OH, which explains the downregulation of the electron transfer module (ETM) and its redox partner (Fig. S10, Table S6). Under “normal” anammox conditions (i.e., NO_2_^−^ as electron acceptor), the membrane associated quinol-interacting ETM encoded in the HZS gene cluster, mediate the first half-reaction for N_2_H_4_ synthesis (*66, 78*). The ETM provides three-electrons to the HZS enzymatic complex for NO reduction to NH_2_OH with the help of an electron shuttle (*78, 79*). N_2_H_4_ is further oxidized to N_2_ by HDH (Fig. 4). The four low-potential electrons released from this reaction must be stored until they are transferred to a redox partner and feed the quinone (quinol) pool within the anammoxosome membrane to build up the membrane potential (*80*). Currently, it is not known how electrons are transported over membranes when NO_2_^−^ is the electron acceptor. Understanding the electron flow and electron carriers is an important next step in anammox research. A recent exciting study showing the structure of the HDH (*80*), revealed that HDH can store up to 192 electrons and it is proposed that the appropriate carriers might specifically dock into the enzyme to get the electrons and transport them to the desired acceptor. This will prevent accidental transfer of the low-redox potential electrons to random acceptors. Interestingly, HDH was one of the most upregulated enzymes in our study when the anode was the electron acceptor, which suggests an increased demand of electron storage and transport when the anode is the electron acceptor. In the typical anammox process, quinol oxidation supplies electrons for the reductive steps, thus closing the electron transfer cycle. However, under electrode-dependent anammox process, where there is no NO_2_^−^ and the electrode is the sole electron acceptor, electrons must first pass to the cytoplasm. By accepting the electrons from N_2_H_4_ oxidation, energy would be conserved as reduced quinone and NAD(P)H, which can work as electron carrier in the cytoplasm. This set of reactions are thermodynamically feasible and are done by the Rieske/cytb complexes of anammox bacteria (*66*) (Supplementary materials, Respiratory complexes of anammox bacteria in EET-dependent anammox process).

Several genes encoding for low-molecular-weight mobile carriers dissolved in the cytoplasm (NADH, ferredoxins, rubredoxins) were found expressed under set potential conditions (Fig. S10, Table S3). These low-molecular-weight electron carriers act as electron shuttles between the respiratory complexes in the anammoxosome and the central carbon and iron metabolism of anammox bacteria (*65, 66*) (Supplementary materials). Even though non-heme electron carriers dissolved in the cytoplasm are proposed as electron shuttles, it is still not clear how the electrons are transferred from the respiratory complexes in the anammoxosome to the inner membrane, even when NO_2_^-^ is the electron acceptor. Our transcriptomic data identified an EET pathway in *Ca*. Brocadia electricigens in response to the electrode as electron acceptor. The EET pathway found in anammox bacteria is analog to the ones present in metal-reducing organisms such as *Geobacter* spp. and *Shewanella* spp (*25*). To overcome the membrane barriers, electrons from the oxidation of menaquinol by an inner-membrane tetraheme *c*-type cytochrome (Cyt *c* (4 hemes)) are transferred to the periplasmic mono-heme *c*-type cytochrome (Cyt *c* (1 heme)) (Fig. S10, Table S3). The tetraheme *c*-type cytochrome (Cyt *c* (4 hemes)), may function as a quinol dehydrogenase of the EET cascade, similar to the role played by the tetraheme CymA in *Shewanella* (*81, 82*). The highly upregulated mono-heme cytochrome c (Cyt *c* (1 heme)) was found to have homology with MtoD of the metal-oxidizing bacteria *Sideroxydans lithotrophicus* ES-1 (*83*). MtoD has been characterized as a periplasmic monoheme cytochrome *c* that works as electron shuttle between CymA and outer membrane cytochromes (*81, 83*). It is still not clear which protein(s) feed the menaquinol pool used by the tetraheme *c*-type cytochrome in the inner membrane. It has been reported that for EET in *S. oneidensis*, electrons could enter the inner-membrane pool via the activity of primary dehydrogenases, such as NADH dehydrogenases, hydrogenases or formate dehydrogenase (Fdh) (*84*). Also, a previous study revealed formate oxidation coupled with Fe(III) or Mn(IV) reduction in anammox bacteria (*6*). In our analysis, we found a significant expression under set potential of multiple copies of the Fdh and its transcriptional activator (Fig. S10, Table S3), which possibly are involved in the EET pathway to respire insoluble minerals in anammox bacteria. It has been reported that C1 metabolism such as formate oxidation by Fdh is strongly related to the electron transferring to the extracellular environment (*84*). Evidence suggests that formate can act as a stimulus for external electron transfer in the absence of soluble electron acceptors, which is related to the existence of a periplasmic Fdh to convert formate to CO_2_ with the electrons being released extracellularly (*84*). Similar to *S. oneidensis*, *Ca*. Brocadia electricigens gets a significant amount of proton motive force and feeds the quinol pool in the inner membrane by transporting and oxidizing formate in the periplasm (*85*) (Fig. S10).

Outer membrane protein complexes can transfer the electrons from the periplasm to the bacterial surface via an electron transport chain (*81*). The wide windows of these cytochromes allow an overlapping of redox potentials in an electron transport chain and make possible a thermodynamic downhill process of electron transport (*86*). It has been reported that *K. stuttgartiensis* possesses a trans-outer membrane porin-cytochrome complex for extracellular electron transfer that is widespread in different phyla (*87, 88*). The genes encoding for the porin-cytochromes are adjacent to each other in the genome (kuste4024 and kuste4025) and consist of a periplasmic and a porin-like *c*-type outer-membrane cytochrome (*87, 88*). As expected, *Ca*. Brocadia electricigens expressed the ortholog of the outer-membrane porin-cytochrome complex (Fig. S10, Table S3). Compared to the porin-cytochrome complexes of six different phyla, anammox bacteria porin-cytochromes are larger and possess more heme-binding motifs (*88*). This may provide anammox bacteria a sufficient span to transfer electrons across the outer membrane without the need of additional outer-membrane cytochromes (*88*). However, biofilm CV analysis (Fig. 2E, Fig. S3C and D) exhibited oxidation/reduction peaks, which suggests that additional cytochrome(s) that transfer electrons directly to the electrode via solvent exposed hemes may be involved. Also, no cytochromes for long-range electron transport were detected in the analysis (Table S6 and 7), suggesting that EET to electrodes by anammox bacteria rely on a direct EET mechanism. Homology detection and structure prediction by hidden Markov model comparison (HMM-HMM) of the highly upregulated penta-heme cytochrome EX330_07910 (Fig. S10, Table S6) gave high probability hits to proteins associated to the extracellular matrix and outer membrane iron respiratory proteins such as MtrF, OmcA and MtrC. Also, it is worth mentioning that the gene cluster EX330_07910-07915 was one of the most upregulated under set-potential conditions. Therefore, future work should focus on determining the role of EX330_07910-07915 in the EET-dependent anammox process. Likewise, we also found the expression of outer membrane mono-heme *c*-type cytochromes (OM Cyt c (1 heme) (Fig. S10, Table S7) homologs to *G. sulfurreducens’* OmcF, which has been characterized to be an outer membrane-associated monoheme cytochrome involved in the regulation of extracellular reduction of metal oxides (*89*). Anammox bacteria have a diverse repertoire of conductive and electron-carrier molecules that can be involved in the electron transfer to insoluble electron acceptors. Therefore, it is possible that different pathways may be involved in parallel in the EET-dependent anaerobic ammonium oxidation

#### Respiratory complexes of anammox bacteria in EET-dependent anammox process

In the current proposed model of the anammox process, the four electrons released from the N_2_H_4_ oxidation are transferred to the menaquinone pool in the anammoxosome membrane by the action of a yet unknown oxidoreductase (*66, 80*). The resulting proton gradient across the anammoxosome membrane drive the adenosine 5′-triphosphate (ATP) synthesis (*80*). However, in general, little is known about how anammox bacteria transport and utilize the energy released by the N_2_H_4_ oxidation in the respiratory complexes in the anammoxosome membrane (*80*). These respiratory processes depend heavily on membrane-bound complexes such as the *bc1* complex (*65*). It is proposed that the Rieske/cytb *bc1* complex in anammox bacteria plays a central role coupling the oxidation of two-electron carrier quinol with the reduction of two *c*-type cytochromes with a net proton translocation stoichiometry of 4H^+^/2e^−^ (*65*). With this electron bifurcation mechanism, it is thermodynamically feasible to synthesize NAD(P)H by coupling oxidation of (mena)quinol to the reduction of an electron acceptor of higher redox potential such as NAD(P) (*65*). A previous study revealed that in the typical anammox process (i.e., NO_2_^−^ as electron acceptor), gene products of Rieske/cytb *bc1* and *bc3* of anammox bacteria were the least and most abundant complexes in the anammoxosome membrane, respectively (*66*). In contrast, our comparative transcriptomics analysis revealed that with electrode as the sole electron acceptor, complex *bc1* and *bc3* were upregulated and downregulated, respectively (Fig. S10, Table S6). In agreement with the current knowledge of anammox biochemistry (*65, 66*), in our model, we also propose a bifurcation mechanism for NAD(P)H generation in concert with menaquinol oxidation catalyzed by the *bc1* complex and/or a H^+^ translocating NADH:quinone oxidoreductase (NADH dehydrogenase, NADH-DH). Energy released by NADH oxidation to quinone reduction (Δ*G*^0′^ = - 47 kJ) can be utilized by the upregulated sodium-dependent NADH:ubiquinone oxidoreductase (RnfABCDGE type electron transport complex) to translocate sodium ions, thus creating a Na-motive force (*65*) (Fig. S10). Accordingly, a Na-motive force might be employed to drive the opposite unfavorable NAD^+^ reduction by the upregulated NAD-dependent oxidoreductases, quinol dehydrogenases or NAD-dependent dehydrogenase (*65*) (Fig. S10, Table S3). In the membrane-bound Rnf complex, the electrons from the oxidation of NADH are transferred to ferredoxins (Fd_red_) (*66*). Since redox potential of Fd (*E*^0′^_Fd_ = −500 to −420 mV) is more negative than NAD+/NADH couple (*E*^0′^_NADH_ = −320 mV), the excess energy is available for transmembrane ion transport (*86*). Ferredoxins act as non-heme electron carriers in the cytoplasm for reactions of the central carbon and iron metabolism of anammox bacteria (*65, 66*).

#### Central carbon metabolism of anammox bacteria in EET-dependent anammox process

Our analysis showed upregulation under electrode-dependent anammox process of the genes in the Wood-Ljungdahl pathway for CO_2_ fixation and acetyl-CoA synthesis (Fig. S10, Table S6). Also, the key enzyme for CO_2_ fixation via the reductive tricarboxylic acid cycle (rTCA) pyruvate:ferredoxin oxidoreductase (PFdO) was upregulated under electrode-dependent anammox process (Fig. S10, Table S6). This enzyme can catalyze the decarboxylation of pyruvate with use of ferredoxins (*90*). Apart from serving as main electron donor in anammox bacteria, NH_4_^+^ is also assimilated for biosynthesis via glutamate synthase (GltS). Multiple copies of GltS were found expressed in our analysis (Fig. S10, Table S3). GltS catalyzes the binding of the ammonium-nitrogen to 2-oxoglutarate with the oxidation of Fd_red_ (*91*). The 2-oxoglutarate used for this reaction can be provided by the key enzyme of the rTCA cycle 2-oxoglutarate:ferredoxin oxidoreductase (OGOR) (*92*). Multiple copies of OGOR were expressed similarly under both types of electron acceptor (Fig. S10, Table S3). These enzymes depend on the reducing power of reduced ferredoxin (Fd_red_) for the reactions, which are the proposed soluble electron carriers in the cytoplasm.

#### Iron assimilation in anammox bacteria in EET-dependent anammox process

Iron is the fourth most abundant element in Earth’s crust (*93*) and plays an essential role in anammox metabolism. Energy conservation in anammox bacteria depends on iron-containing proteins (i.e., cytochromes and iron-sulfur proteins) (*14*). Surprisingly the proteins involved in iron transport and assimilation are still unknown. Our analysis revealed that in the absence of soluble electron acceptors (i.e., NO_2_^-^, NO_3_^-^), *Ca*. Brocadia electricigens expressed two gene clusters encoding a siderophore-mediated iron uptake system (Fig. S10, Table S3 and 13). The expressed siderophore-mediated transport system, which was previously believed to be absent in anammox bacteria (*14*), is homolog to the well-studied TonB-dependent Fe(III) uptake complex present in Gram-negative bacteria (*94*). Fe(III) uptake relies on beta-barrel TonB-dependent receptors in the outer membrane (*95*) and an energy-transducing protein complex TonB-ExbB-ExbD that links the outer with the inner membrane and generate a proton motive force (*94*). A periplasmic iron-binding protein and an ATP-dependent ABC transporter permease are responsible for the Fe(III)-siderophore translocation across the inner membrane into the cytoplasm, where the Fe(III) is reduced to Fe(II) and released from the complex (*94*) (Fig. S10). Fe(III) reduction in the cytoplasm can be carried out by ferric-chelate reductases/rubredoxins, from which multiple genes were found to be expressed (Fig. S10, Table S3). After being reduced, the iron can be assimilated into the metalloprosthetic groups of protein complexes (*14*). Even though Fe(III) was not added in the experimental setup, *Ca*. Brocadia electricigens may have activated this system in order to uptake Fe(III) as an alternative electron acceptor as well as for iron uptake for assimilation. This finding is in agreement with a previous study using the EET-capable model bacteria *Geobacter sulfurreducens* (*8*)*, in which it was shown that t*he pathways required for EET and metal oxide reduction are distinct.

**Figure S1.**
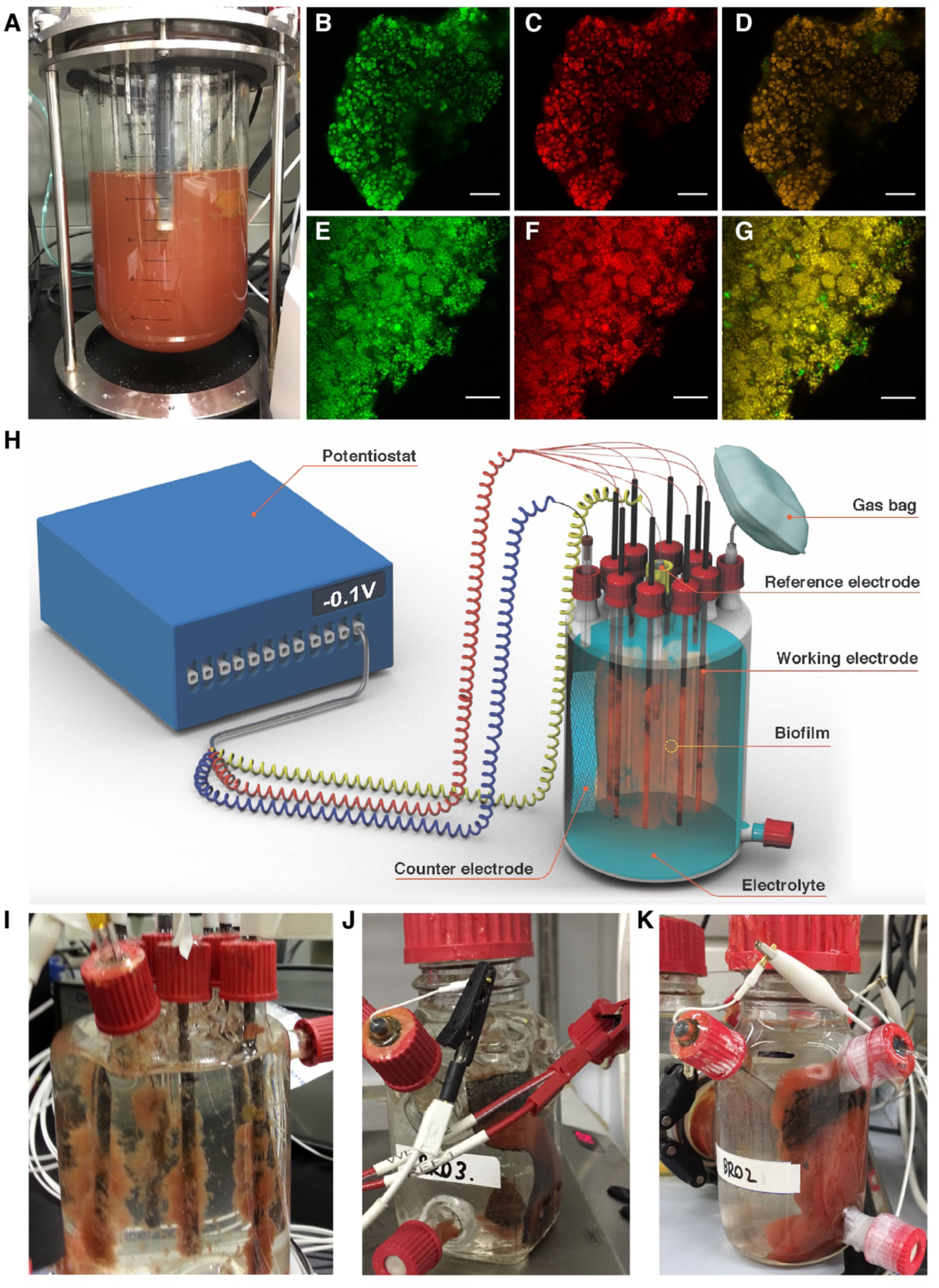
Reactors used in this study. (**A**) Photograph of membrane bioreactor (MBR) used for the enrichment of anammox planktonic cells. (**B** to **G**) Confocal laser scanning microscopy images of enriched biomass of *Ca*. Brocadia (**B**, **C** and **D**) and *Ca*. Scalindua (**E**, **F** and **G**). The images are showing all bacteria (green), anammox bacteria (red) and the merged micrograph (yellow). Fluorescence *in-situ* hybridization was performed with EUB I, II and III probes for all bacteria and Alexa647-labeled Amx820 probe for anammox bacteria. The scale bars represent 20 μm in length. (**H**) Schematic representation of the multiple working electrode microbial electrolysis cell (MEC). (**I** to **K**) Photographs of the single-chamber multiple working electrode MEC (**I**); single-chamber MEC with single working electrode (**J**); and double-chamber MEC with single working electrode (**K**) with anammox biofilm shown on all the electrodes.

**Figure S2.**
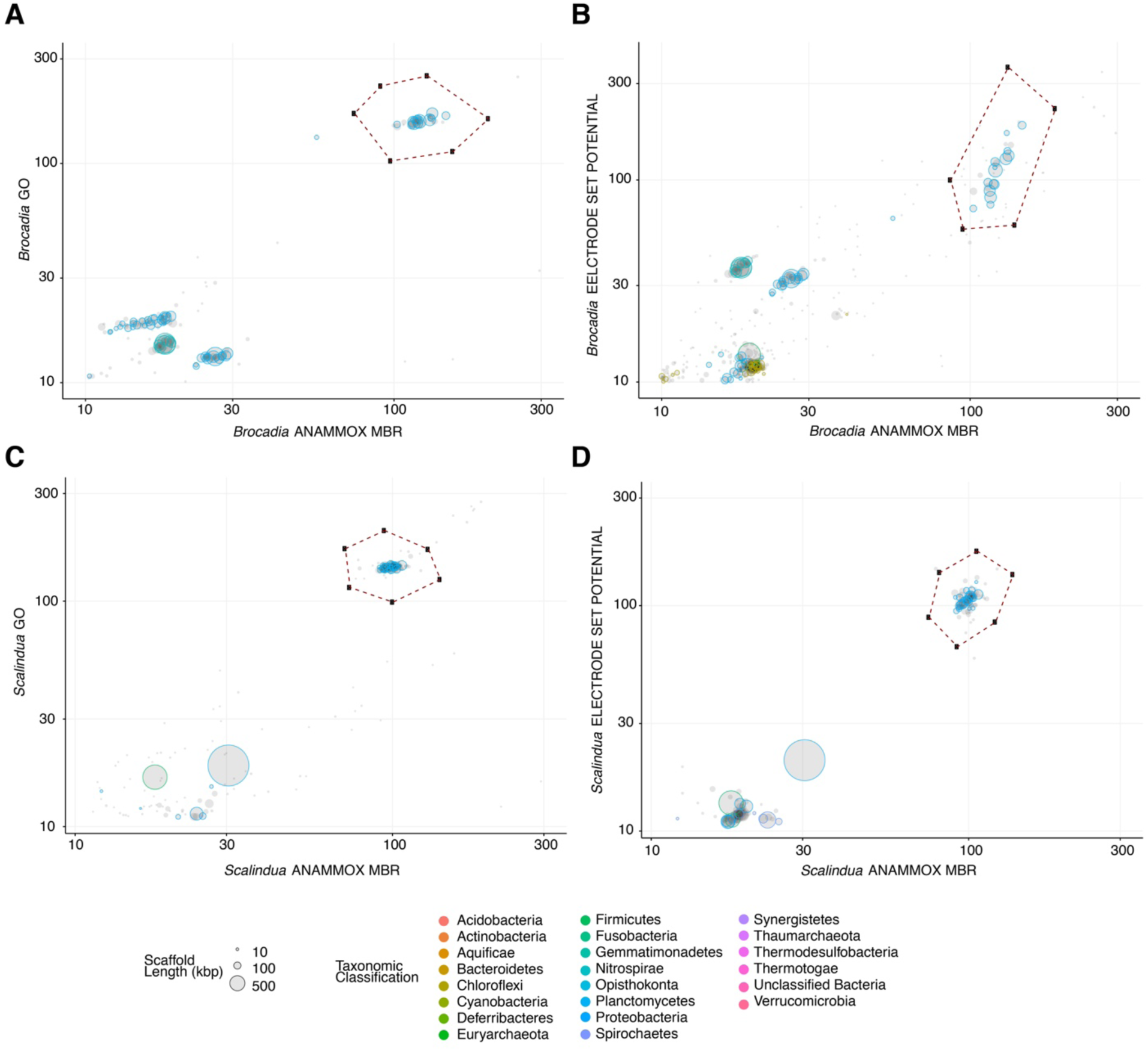
Sequence composition-independent binning of the metagenome scaffolds from graphene oxide (GO) and microbial electrolysis cell (MEC) experiments. (**A** to **D**) Differential coverage binning of the genome sequences from the incubation of *Ca*. Brocadia and *Ca*. Scalindua with GO (**A** and **C**) or working electrode (0.4 V vs Ag/AgCl applied potential) as the sole electron acceptor (**B** and **D**). Each circle represents a metagenomic scaffold, with size proportional to scaffold length; only scaffolds ≥ 5 Kbp are shown. Taxonomic classification is indicated by color; clusters of similarly colored circles represent potential genome bins. The x and y-axes show the sequencing coverage in the samples (log-scaled). Extracted anammox genomes are enclosed by dashed polygons.

**Figure S3.**
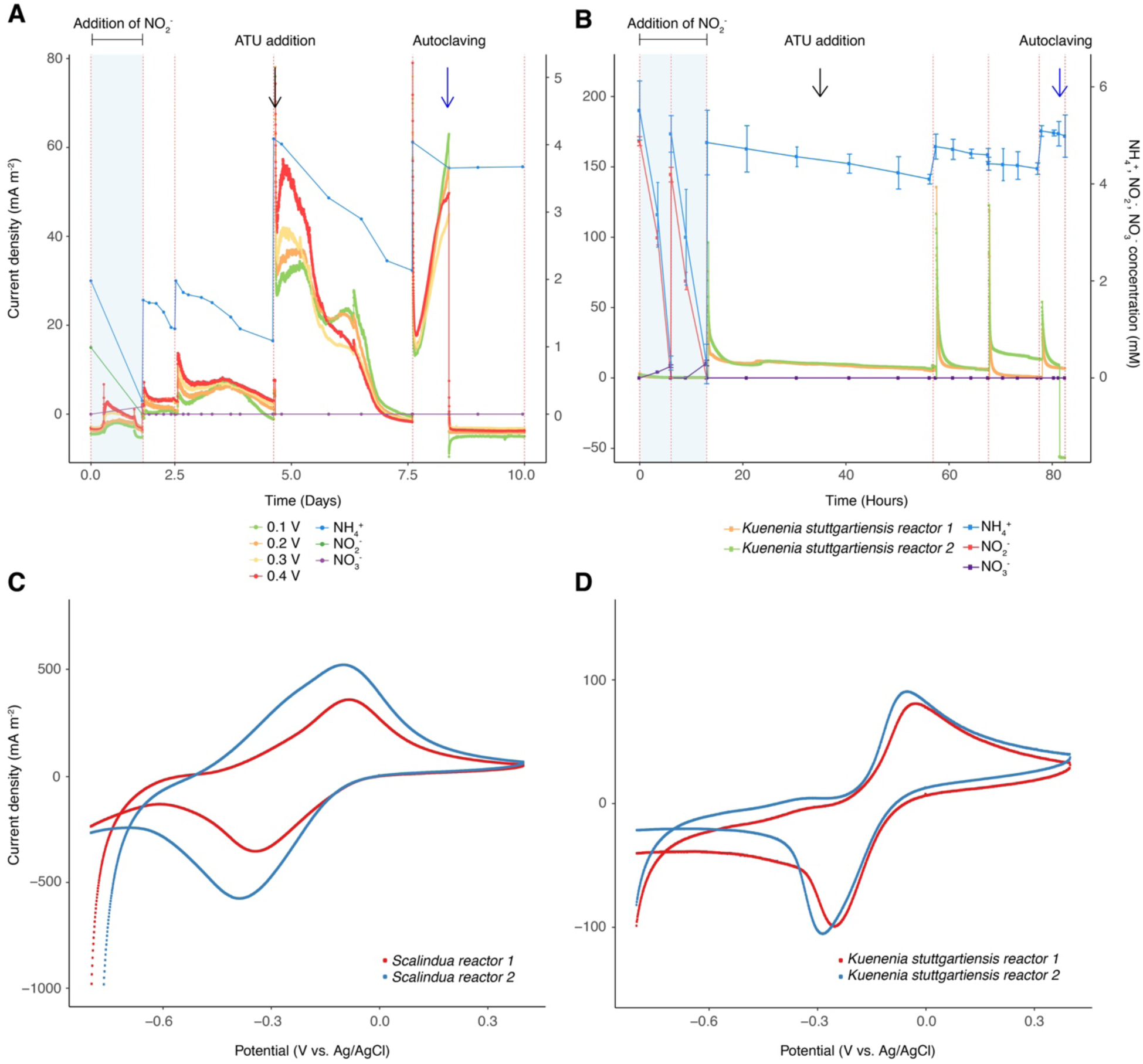
*Ca.* Scalindua and *Kuenenia stuttgartiensis* are electrochemically active. (**A** and **B**) Ammonium oxidation coupled to current generation in chronoamperometry experiment conducted in (**A**) single-chamber multiple working electrode MEC inoculated with *Ca*. Scalindua and operated under different set potentials and (**B**) single-chamber MECs inoculated with *Kuenenia stuttgartiensis* and operated with a working electrode at 0.4 V vs Ag/AgCl. Red dashed lines represent a change of batch. The highlighted area with blue refers to the operation of MEC in the presence of nitrite, which is the preferred electron acceptor for anammox bacteria. The black arrow indicates addition of *allylthiourea* (ATU), a compound that selectively inhibits nitrifiers. The black arrow in plot (**B**) indicates ATU addition in reactor 2 of *Kuenenia stuttgartiensis*. The blue arrow indicates autoclaving followed by re-connecting of the MECs. The blue arrow in plot (**B**) indicates autoclaving of reactor 2 of *Kuenenia stuttgartiensis*. (**C** and **D**) Cyclic Voltammogram (1 mV s^-1^) of *Ca*. Scalindua (**C**) and *Kuenenia stuttgartiensis* (**D**) biofilm grown on anode.

**Figure S4.**
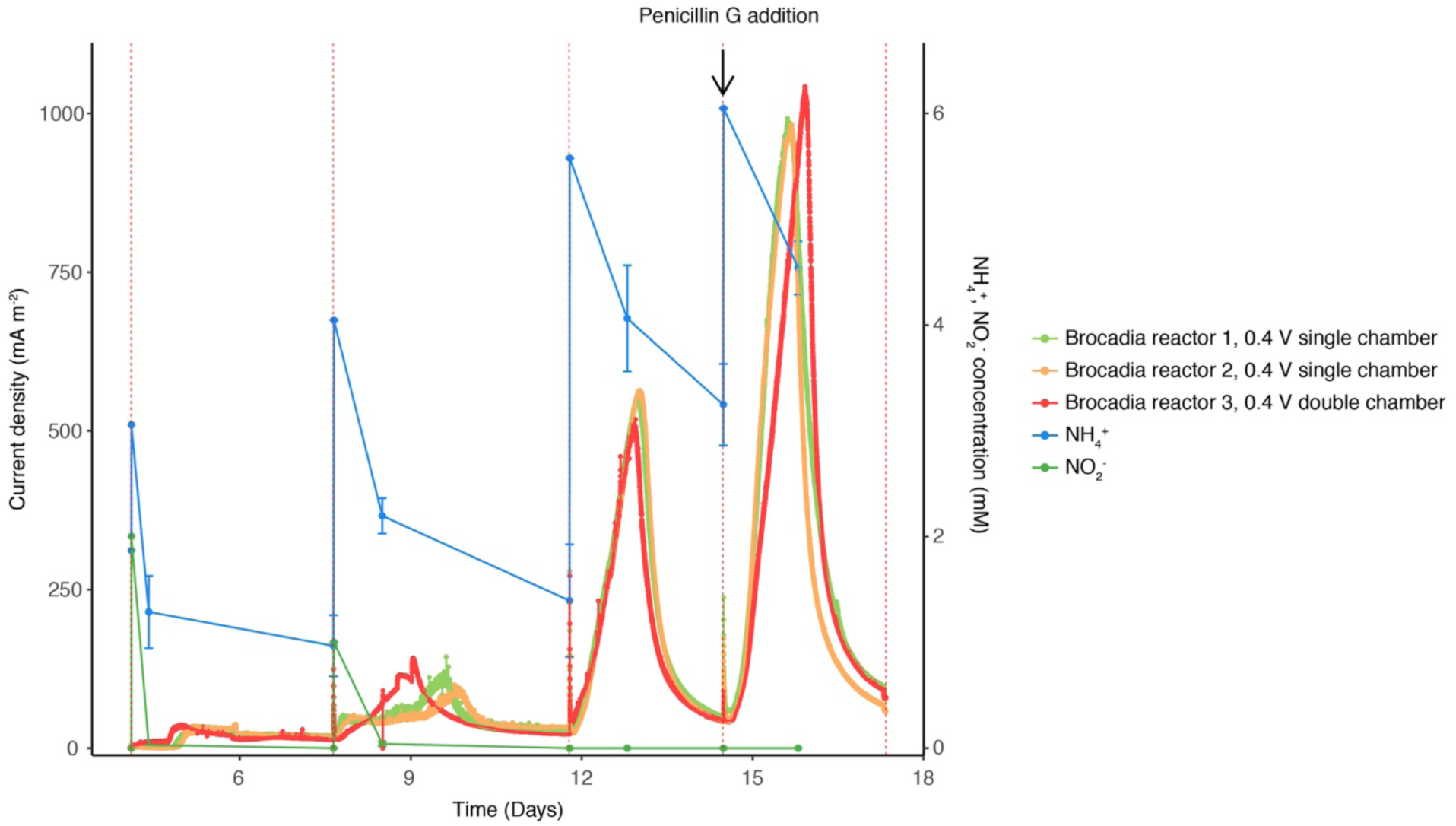
Influence of cathodic reaction (i.e., hydrogen evolution reaction) on electrode-dependent anaerobic ammonium oxidation by *Ca*. Brocadia. Ammonium oxidation and chronoamperometry of single and double-chamber MECs inoculated with *Ca*. Brocadia and operated at set potential of 0.4 V vs Ag/AgCl. Red dashed lines represent a change of batch. The black arrow indicates addition of penicillin G to reactor 2. Penicillin G is not active against anammox bacteria but inhibits the activity of heterotrophs.

**Figure S5.**
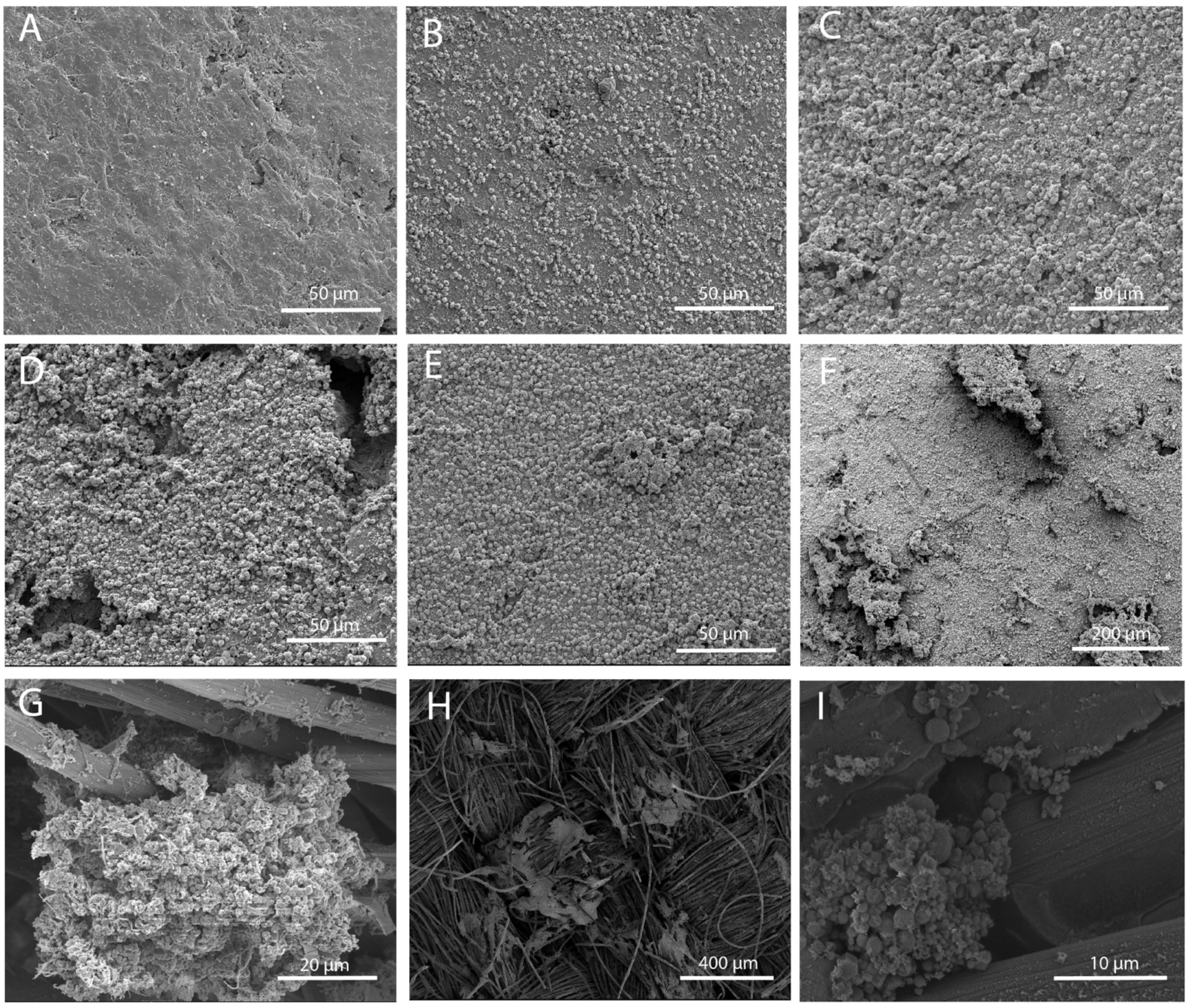
Micrographs of anammox biofilm on working electrodes. (**A** to **F**) Scanning electron microscopy (SEM) images of *Ca*. Brocadia biofilm grown on graphite rod anodes after 55 days of operation at set potential of 0 V (**A**), 0.1 V (**B**), 0.2 V (**C**), 0.3 V (**D**) and 0.4 V (**E** and **F**) vs Ag/AgCl. (**G**) SEM image showing *Ca*. Scalindua biofilm grown on carbon cloth anode at set potential of 0.4 V vs Ag/AgCl. (**H** and **I**) SEM images of *Kuenenia stuttgartiensis* biofilm grown on carbon cloth anode at 0.4 V vs Ag/AgCl.

**Figure S6.**
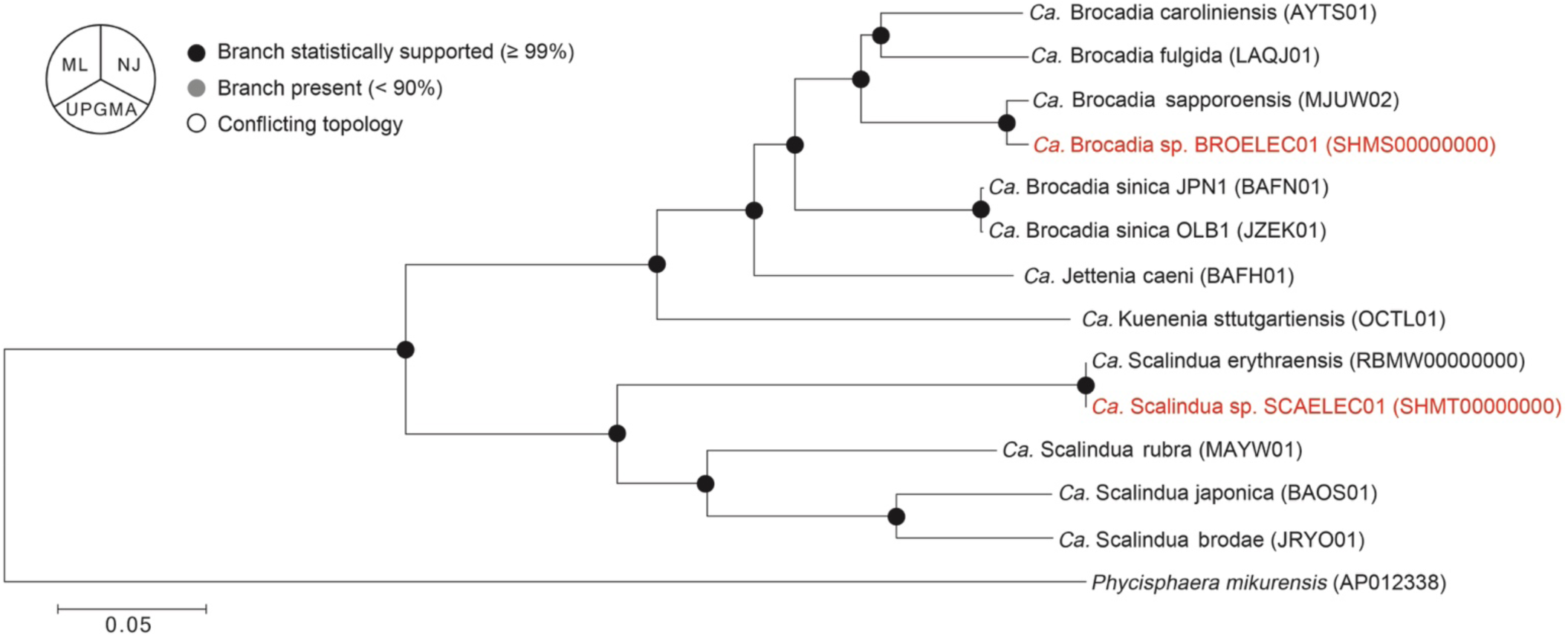
Phylogenomics analysis of anammox genomes extracted from the working electrodes of *Ca*. Brocadia and *Ca*. Scalindua MECs and closely related genomes downloaded from the NCBI genome repository. Pie charts at the nodes represent the bootstrap support values and the bootstrap consensus inferred from 1000 iterations. Support value ≥ 99% is filled with black. ML, maximum likelihood method; NJ, neighbor joining method; and UPGMA, unweighted pair group method with arithmetic mean. The anammox genomes extracted from the biofilm community on the working electrodes are shown in red. GenBank accession numbers for each genome are provided in parentheses. Sequences of two different strains of *Brocadia sinica* were used as a reference of same species genomes. Sequence of a member from the phylum planctomycetes different than anammox bacteria was used as outgroup.

**Figure S7.**
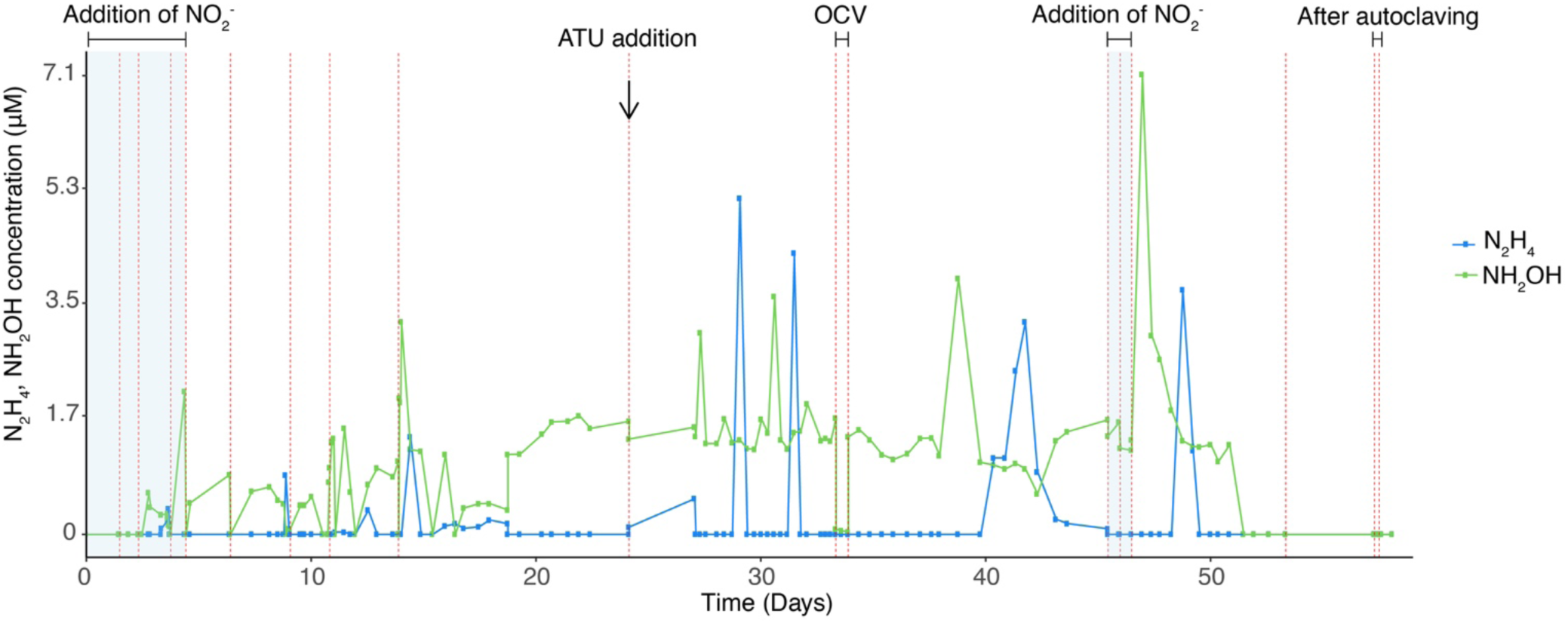
Time course of the concentration of hydroxylamine (NH_2_OH) and hydrazine (N_2_H_4_) in chronoamperometry experiment conducted in single-chamber multiple working electrode MEC inoculated with *Ca.* Brocadia and operated under different set potentials. Red dashed lines represent a change of batch. The highlighted area in blue refers to the operation of MEC in the presence of nitrite, which is the preferred electron acceptor for anammox bacteria. The black arrow indicates addition of ATU. OCV indicates MEC operated under open circuit voltage.

**Figure S8.**
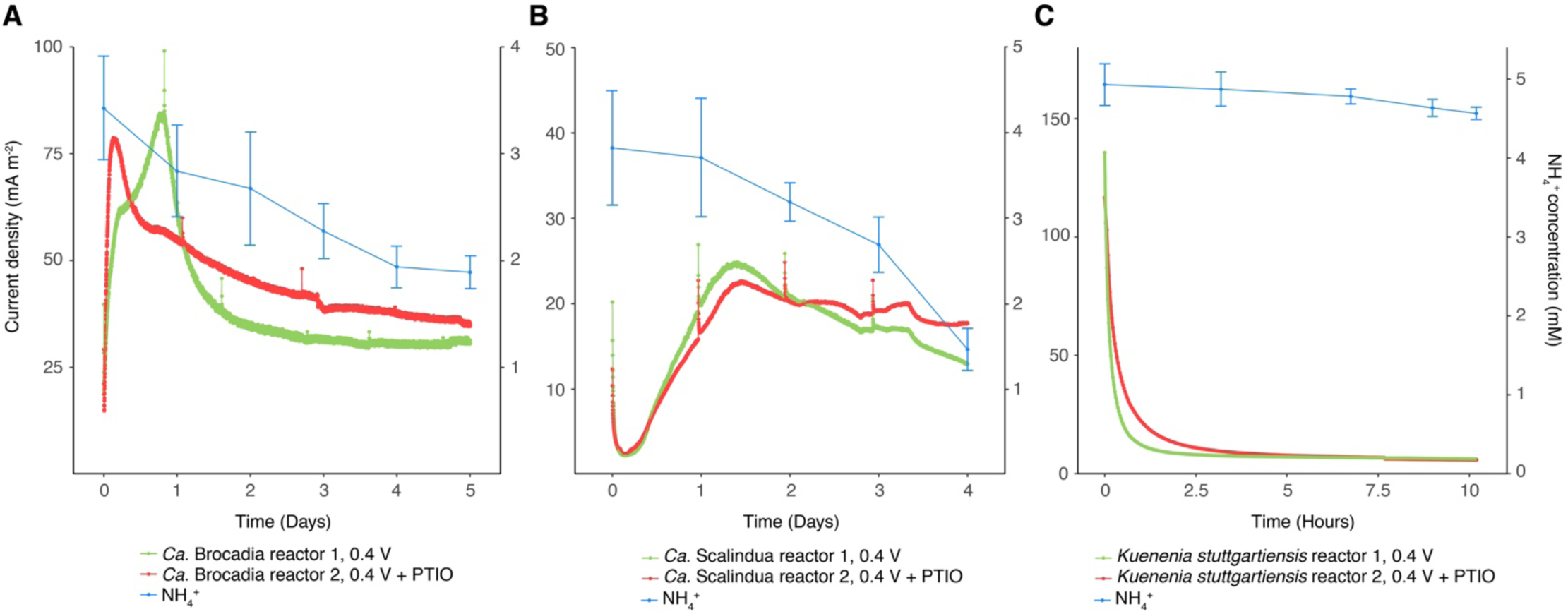
Influence of PTIO, a NO-scavenger, on electrode-dependent anaerobic ammonium oxidation by different anammox bacteria. (**A**) Ammonium oxidation and chronoamperometry of *Ca*. Brocadia single-chamber MECs with and without PTIO addition. (**B**) Ammonium oxidation and chronoamperometry of *Ca*. Scalindua single-chamber MECs with and without PTIO addition. (**C**) Ammonium oxidation and chronoamperometry of *Kuenenia stuttgartiensis* single-chamber MECs with and without PTIO addition.

**Figure S9.**
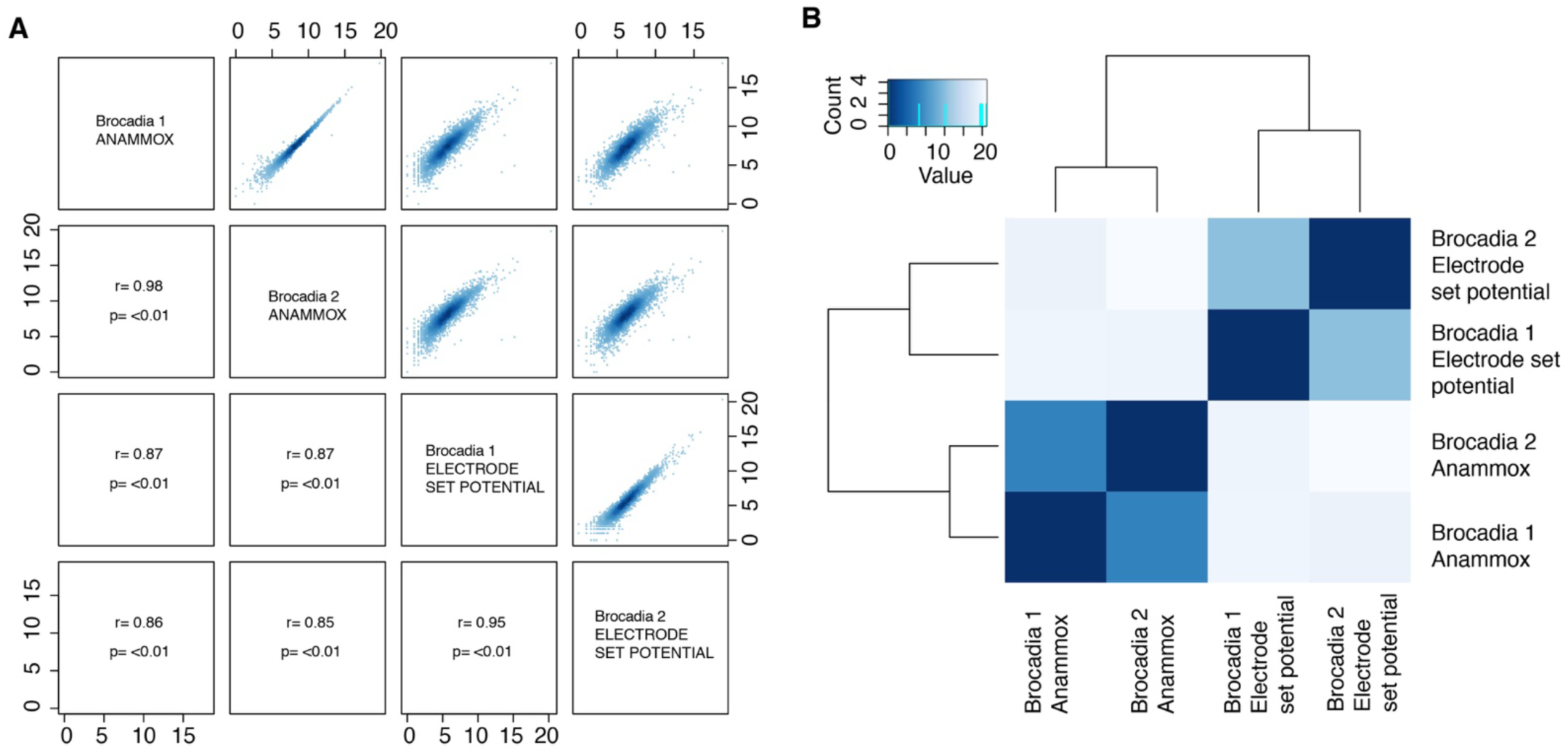
Overall transcriptomics similarity between biological replicate samples. (**A**) Pairwise overview of *Brocadia* transcriptomics samples. “Brocadia Anammox” corresponds to the experimental condition where nitrite was used as the sole electron acceptor. “Brocadia Electrode set potential” corresponds to the experimental condition where the working electrode (0.4 V vs Ag/AgCl) was used as the sole electron acceptor. All counts were normalized to Log2+1 values. The upper right panel shows the normalized counts while the lower-left panel shows the corresponding Pearson correlation coefficient between samples and the P-value. The intra-replicate correlation is high and conversely the inter-replicate correlation is low. This indicates high similarity between the biological replicates and differentially expressed genes across the experimental setups. (**B**) Heatmap clustering of the sample-to-sample distances. Hierarchical clustering shows that anammox (nitrite as electron acceptor) and electrode-dependent anammox (set potential of 0.4 V vs Ag/AgCl) samples were clearly separated and clustered as independent groups.

**Figure S10.**
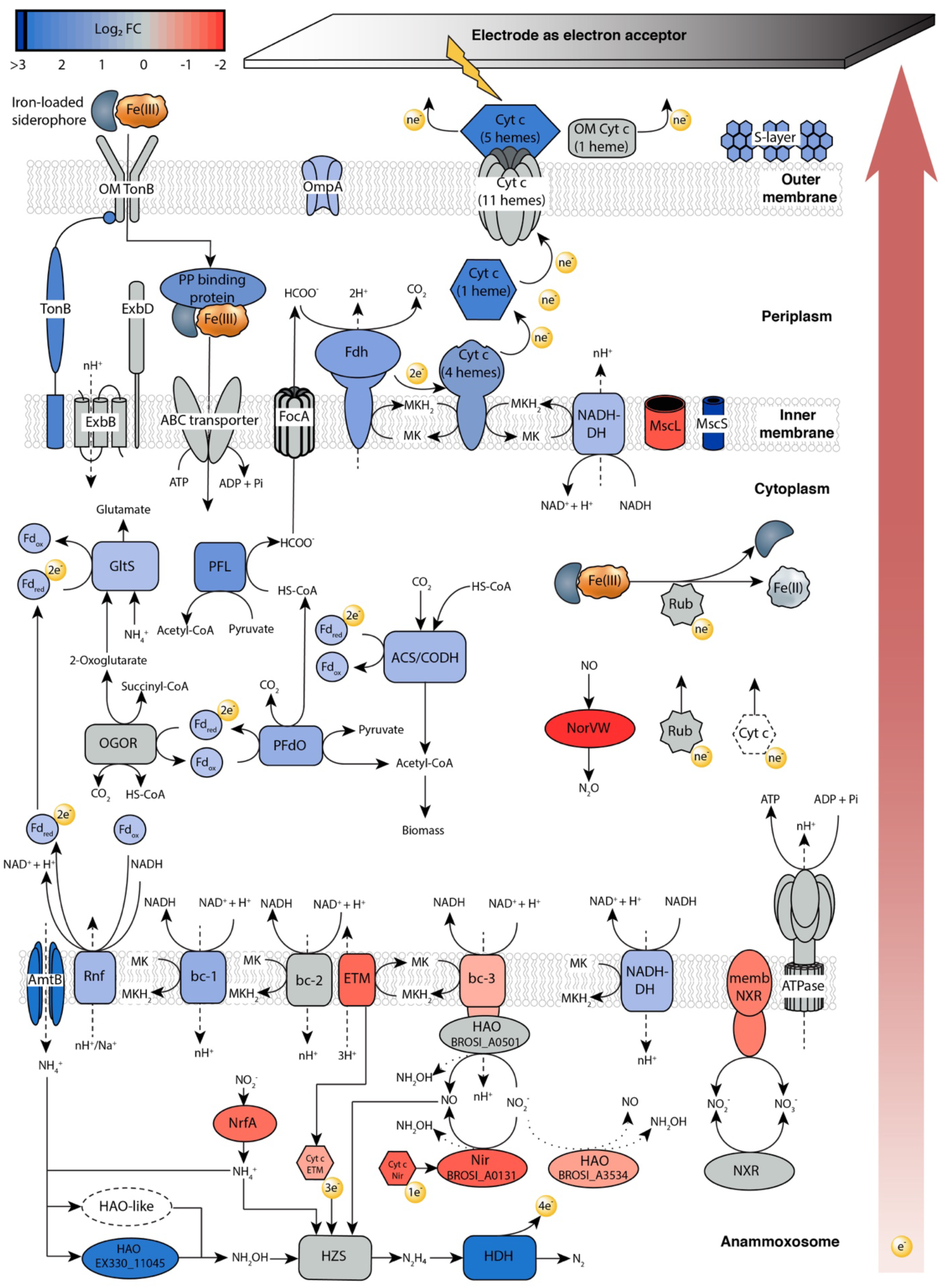
Molecular model of electrode-dependent anaerobic ammonium oxidation. The putative EET metabolic pathway of *Ca.* Brocadia to deliver electrons to an electrode was constructed with the transcriptional changes of selected marker genes in response to the electrode as the electron acceptor. Samples for comparative transcriptomic analysis were taken from mature electrode’s biofilm of single-chamber MECs with NO_2_^-^ as sole electron acceptor and after switching to set potential growth (0.4 V vs Ag/AgCl, electrode as electron acceptor). Log_2_ fold changes (Log_2_ FC) in expression are shown as follows; gene and gene clusters are shown in blue if upregulated or red if downregulated relative to the electrode as the electron acceptor. The color grey corresponds to genes that were expressed under similar levels in both conditions (i.e., electrode or nitrite as electron acceptor). Dashed lines represent proton (H^+^) transport across membranes. Dashed curves indicate proteins, reactions or processes that have not been established yet. Genomic identifiers of the proteins used in the model are listed in Table S5. The reactions are described in the Supplementary Text of the paper and are catalyzed by the enzymes ABC transporter: Iron ABC transporter permease and ATP-binding protein; ACS/CODH: Acetyl-CoA synthase/ CO dehydrogenase; AmtB: Ammonium transport protein; ATPase: ATP synthase; bc-1: Rieske/cytochrome b complex; bc-2: Rieske/cytochrome b complex; bc-3: Rieske/cytochrome b complex; Cyt *c* (1 heme): Periplasmic mono-heme *c*-type cytochrome; Cyt *c* (11 hemes): Membrane-anchored undeca-heme cytochrome *c*; Cyt *c* (4 hemes): Membrane-anchored tetraheme *c*-type cytochrome; Cyt *c* (5 hemes): Outer membrane penta-heme *c*-type cytochrome; Cyt *c* ETM: Cytochrome c redox partner of the ETM; Cyt Nir: Cytochrome *c*; ETM: electron transfer module for hydrazine synthesis; ExbB: Biopolymer transport protein ExbB/TolQ; ExbD: Biopolymer transport protein ExbD/TolR; FDH: membrane-bound formate dehydrogenase; Fd_ox_: Ferredoxin (oxidized); Fd_red_ Ferredoxin (reduced); FocA: Formate/nitrite transporter; GltS: Glutamate synthase; HAO BROSI_A0501: Hydroxylamine oxidoreductase; HAO BROSI_A3534: Hydroxylamine oxidoreductase; HAO EX330_11045: Hydroxylamine oxidoreductase; HDH: hydrazine dehydrogenase; HZS: hydrazine synthase; membNXR: membrane-bound complex of the *nxr* gene cluster; MscL: Large mechanosensitive channel; MscS: Pore-forming small mechanosensitive channel; NADH-DH: NADH dehydrogenase; Nir BROSI_A0131: nitrite reductase; NorVW: Flavodoxin nitric oxide reductase; NrfA: ammonium-forming nitrite reductase; NXR: nitrite:nitrate oxidoreductase; OGOR: 2-oxoglutarate ferredoxin oxidoreductase; OM Cyt *c* (1 heme): Outer membrane lipoprotein mono-heme c-type cytochrome; OM TonB: TonB-dependent receptor; OmpA: OmpA-like outer membrane protein, porin; PFdO: Pyruvate ferredoxin oxidoreductase; PFL: Pyruvate formate lyase; PP binding protein: Iron ABC transporter periplasmic substrate-binding protein; Rnf: RnfABCDGE type electron transport complex; Rub: Rubredoxin/ferric-chelate reductase; S-layer: S-layer protein; TonB: Energy transducer TonB.

## References

1. A. H. Devol, Denitrification, Anammox, and N2 Production in Marine Sediments. Ann. Rev. Mar. Sci. 7, 403–423 (2015).

2. P. Lam, M. M. M. Kuypers, Microbial Nitrogen Cycling Processes in Oxygen Minimum Zones. Ann. Rev. Mar. Sci. 3, 317–345 (2010).

3. B. Kartal, J. G. Kuenen, M. C. M. van Loosdrecht, Sewage Treatment with Anammox. Science (80-.). 328, 702–703 (2010).

4. B. Kartal, W. J. Maalcke, N. M. de Almeida, I. Cirpus, J. Gloerich, W. Geerts, H. J. M. Op den Camp, H. R. Harhangi, E. M. Janssen-Megens, K.-J. Francoijs, H. G. Stunnenberg, J. T. Keltjens, M. S. M. Jetten, M. Strous, Molecular mechanism of anaerobic ammonium oxidation. Nature. 479, 127–130 (2011).

5. Z. Hu, H. J. C. T. Wessels, T. van Alen, M. S. M. Jetten, B. Kartal, Nitric oxide-dependent anaerobic ammonium oxidation. Nat. Commun. 10, 1244 (2019).

6. M. Strous, E. Pelletier, S. Mangenot, T. Rattei, A. Lehner, M. W. Taylor, N. Fonknechten, M. Horn, H. Daims, D. Bartol-mavel, P. Wincker, C. Schenowitz-truong, C. Me, A. Collingro, D. Vallenet, B. Snel, B. E. Dutilh, H. J. M. O. Den Camp, C. Van Der Drift, I. Cirpus, K. T. Van De Pas-schoonen, H. R. Harhangi, L. Van Niftrik, M. Schmid, J. Keltjens, J. Van De Vossenberg, B. Kartal, H. Meier, D. Frishman, M. A. Huynen, H. Mewes, J. Weissenbach, M. S. M. Jetten, M. Wagner, D. Le Paslier, Deciphering the evolution and metabolism of an anammox bacterium from a community genome. 440, 790–794 (2006).

7. J. Van De Vossenberg, J. E. Rattray, W. Geerts, B. Kartal, L. Van Niftrik, E. G. Van Donselaar, J. S. S. Damsté, M. Strous, M. S. M. Jetten, Enrichment and characterization of marine anammox bacteria associated with global nitrogen gas production. 10, 3120–3129 (2008).

8. F. Jiménez Otero, C. H. Chan, D. R. Bond, Identification of Different Putative Outer Membrane Electron Conduits Necessary for Fe(III) Citrate, Fe(III) Oxide, Mn(IV) Oxide, or Electrode Reduction by Geobacter sulfurreducens. J. Bacteriol. 200, e00347–18 (2018).

9. K. Rabaey, L. Angenent, U. Schröder, J. Keller, Bioelectrochemical Systems: From Extracellular Electron Transfer to Biotechnological Application (2009), p. Chapter 5, doi:10.2166/9781780401621.

10. D. R. Lovley, J. D. Coates, E. L. Blunt-Harris, E. J. P. Phillips, J. C. Woodward, Humic substances as electron acceptors for microbial respiration. Nature. 382, 445–448 (1996).

11. E. E. Rios-Del Toro, E. I. Valenzuela, J. E. Ramírez, N. E. López-Lozano, F. J. Cervantes, Anaerobic Ammonium Oxidation Linked to Microbial Reduction of Natural Organic Matter in Marine Sediments. Environ. Sci. Technol. Lett. 5, 571–577 (2018).

12. C. Koch, F. Harnisch, Is there a Specific Ecological Niche for Electroactive Microorganisms? ChemElectroChem. 3, 1282–1295 (2016).

13. S. H. Light, L. Su, R. Rivera-Lugo, J. A. Cornejo, A. Louie, A. T. Iavarone, C. M. Ajo-Franklin, D. A. Portnoy, A flavin-based extracellular electron transfer mechanism in diverse Gram-positive bacteria. Nature, 1 (2018).

14. C. Ferousi, S. Lindhoud, F. Baymann, B. Kartal, M. S. M. Jetten, J. Reimann, Iron assimilation and utilization in anaerobic ammonium oxidizing bacteria. Curr. Opin. Chem. Biol. 37, 129–136 (2017).

15. M. Oshiki, T. Awata, T. Kindaichi, H. Satoh, S. Okabe, Cultivation of Planktonic Anaerobic Ammonium Oxidation (Anammox) Bacteria Using Membrane Bioreactor. Microbes Environ. 28, 436–443 (2013).

16. E. C. Salas, Z. Sun, A. Lüttge, J. M. Tour, Reduction of Graphene Oxide via Bacterial Respiration. ACS Nano. 4, 4852–4856 (2010).

17. S. Kalathil, K. P. Katuri, A. S. Alazmi, S. Pedireddy, N. Kornienko, P. M. F. J. Costa, P. E. Saikaly, Bioinspired Synthesis of Reduced Graphene Oxide-Wrapped Geobacter sulfurreducens as a Hybrid Electrocatalyst for Efficient Oxygen Evolution Reaction. Chem. Mater. 31, 3686–3693 (2019).

18. T. Lotti, R. Kleerebezem, C. Lubello, M. C. M. van Loosdrecht, Physiological and kinetic characterization of a suspended cell anammox culture. Water Res. 60, 1–14 (2014).

19. M. A. H. J. Van Kessel, D. R. Speth, M. Albertsen, P. H. Nielsen, H. J. M. O. Den Camp, B. Kartal, M. S. M. Jetten, S. Lücker, Complete nitrification by a single microorganism. Nature. 528, 555–559 (2015).

20. E. Marsili, D. B. Baron, I. D. Shikhare, D. Coursolle, J. A. Gralnick, D. R. Bond, Shewanella secretes flavins that mediate extracellular electron transfer. PNAS. 105, 6–11 (2008).

21. M. Oshiki, S. Ishii, K. Yoshida, N. Fujii, M. Ishiguro, H. Satoh, S. Okabe, Nitrate-Dependent Ferrous Iron Oxidation by Anaerobic Ammonium Oxidation (Anammox) Bacteria. Appl. Environ. Microbiol. 79, 4087–4093 (2013).

22. Z. Hu, T. van Alen, M. S. M. Jetten, B. Kartal, Lysozyme and penicillin inhibit the growth of anaerobic ammonium-oxidizing planctomycetes. Appl. Environ. Microbiol. 79, 7763–9 (2013).

23. M. Albertsen, P. Hugenholtz, A. Skarshewski, K. L. Nielsen, G. W. Tyson, P. H. Nielsen, Genome sequences of rare, uncultured bacteria obtained by differential coverage binning of multiple metagenomes. Nat. Biotechnol. 31, 533–538 (2013).

24. W. J. Maalcke, J. Reimann, S. De Vries, J. N. Butt, A. Dietl, N. Kip, U. Mersdorf, T. R. M. Barends, M. S. M. Jetten, J. T. Keltjens, B. Kartal, Characterization of anammox hydrazine dehydrogenase, a Key-producing enzyme in the global nitrogen cycle. J. Biol. Chem. 291, 17077–17092 (2016).

25. A. Kumar, L. H.-H. Hsu, P. Kavanagh, F. Barrière, P. N. L. Lens, L. Lapinsonnière, J. H. Lienhard V, U. Schröder, X. Jiang, D. Leech, The ins and outs of microorganism– electrode electron transfer reactions. Nat. Rev. Chem. 1, 0024 (2017).

25. A. A. van de Graaf, A. Mulder, P. De Bruijn, M. S. Jetten, L. A. Robertson, J. G. Kuenen, Anaerobic oxidation of ammonium is a biologically mediated process. Appl Env. Microb. 61, 1246–1251 (1995).

26. M. Ali, M. Oshiki, T. Awata, K. Isobe, Z. Kimura, H. Yoshikawa, D. Hira, T. Kindaichi, H. Satoh, T. Fujii, S. Okabe, Physiological characterization of anaerobic ammonium oxidizing bacterium “CandidatusJettenia caeni.” Environ. Microbiol. 17, 2172–2189 (2015).

27. S. X. Peng, M. J. Strojnowski, J. K. Hu, B. J. Smith, T. H. Eichhold, K. R. Wehmeyer, S. Pikul, N. G. Almstead, Gas chromatographic-mass spectrometric analysis of hydroxylamine for monitoring the metabolic hydrolysis of metalloprotease inhibitors in rat and human liver microsomes. J. Chromatogr. B Biomed. Sci. Appl. 724, 181–187 (1999).

28. A. D. and E.W., Baird, R.B., Eaton, L. S. (eds) Clesceri, American Public Health Association, American Water Works Association and Water Environment Federation (2012) Standard Methods for the Examination of Water and Wastewater. (American Public Health Association, New York, ed. 22th, 2012).

29. M. Oshiki, M. Ali, K. Shinyako-hata, H. Satoh, S. Okabe, Hydroxylamine-dependent Anaerobic Ammonium Oxidation (Anammox) by “ Candidatus Brocadia sinica”. Env. Microbiol. 18, 3133–3143 (2016).

30. D. S. Frear, R. C. Burrell, Spectrophotometric Method for Determining Hydroxylamine Reductase Activity in Higher Plants. Anal. Chem. 27, 1664–1665 (1955).

31. G. W. Watt, J. D. Chrisp, Spectrophotometric Method for Determination of Hydrazine. Anal. Chem. 24, 2006–2008 (1952).

32. P. H. Nielsen, H. Daims, H. Lemmer, I. Arslan-Alaton, T. Olmez-Hanci, Eds., FISH Handbook for Biological Wastewater Treatment (IWA Publishing, 2009).

33. H. Daims, A. Brühl, R. Amann, K.-H. Schleifer, M. Wagner, The Domain-specific Probe EUB338 is Insufficient for the Detection of all Bacteria: Development and Evaluation of a more Comprehensive Probe Set. Syst. Appl. Microbiol. 22, 434–444 (1999).

34. R. I. Amann, B. J. Binder, R. J. Olson, S. W. Chisholm, R. Devereux, D. A. Stahl, Appl. Environ. Microbiol., in press (available at http://aem.asm.org/content/56/6/1919.abstract).

35. M. Schmid, U. Twachtmann, M. Klein, M. Strous, S. Juretschko, M. Jetten, J. W. Metzger, K.-H. Schleifer, M. Wagner, Molecular Evidence for Genus Level Diversity of Bacteria Capable of Catalyzing Anaerobic Ammonium Oxidation. Syst. Appl. Microbiol. 23, 93–106 (2000).

36. M. Schmid, K. Walsh, R. Webb, W. I. Rijpstra, K. van de Pas-Schoonen, M. J. Verbruggen, T. Hill, B. Moffett, J. Fuerst, S. Schouten, J. S. Sinninghe Damsté, J. Harris, P. Shaw, M. Jetten, M. Strous, Candidatus “Scalindua brodae”, sp. nov., Candidatus “Scalindua wagneri”, sp. nov., Two New Species of Anaerobic Ammonium Oxidizing Bacteria. Syst. Appl. Microbiol. 26, 529–538 (2003).

37. H. Daims, Use of Fluorescence In Situ Hybridization and the daime Image Analysis Program for the Cultivation-Independent Quantification of Microorganisms in Environmental and Medical Samples. 4, 1–8 (2009).

38. M. F. Alqahtani, K. P. Katuri, S. Bajracharya, Y. Yu, Z. Lai, P. E. Saikaly, Porous Hollow Fiber Nickel Electrodes for Effective Supply and Reduction of Carbon Dioxide to Methane through Microbial Electrosynthesis. Adv. Funct. Mater. 28, 1–8 (2018).

39. P. Walther, A. Ziegler, Freeze substitution of high-pressure frozen samples : the visibility of biological membranes is improved when the substitution. J. Microsc. 208, 3–10 (2002).

40. M. Martin, Cutadapt removes adapter sequences from high-throughput sequencing reads. EMBnet.journal. 17, 10–12 (2011).

41. A. Bankevich, S. Nurk, D. Antipov, A. A. Gurevich, M. Dvorkin, A. S. Kulikov, V. M. Lesin, S. I. Nikolenko, S. Pham, A. D. Prjibelski, A. V. Pyshkin, A. V. Sirotkin, N. Vyahhi, G. Tesler, M. a. Alekseyev, P. a. Pevzner, SPAdes: A New Genome Assembly Algorithm and Its Applications to Single-Cell Sequencing. J. Comput. Biol. 19, 455–477 (2012).

42. H. Li, Minimap2: pairwise alignment for nucleotide sequences. Bioinformatics. 34, 3094–3100 (2018).

43. H. Li, B. Handsaker, A. Wysoker, T. Fennell, J. Ruan, N. Homer, G. Marth, G. Abecasis, R. Durbin, The Sequence Alignment/Map format and SAMtools. Bioinformatics. 25, 2078–2079 (2009).

44. D. Hyatt, G.-L. Chen, P. F. LoCascio, M. L. Land, F. W. Larimer, L. J. Hauser, Prodigal: prokaryotic gene recognition and translation initiation site identification. BMC Bioinformatics. 11, 119 (2010).

45. C. L. Dupont, D. B. Rusch, S. Yooseph, M.-J. Lombardo, R. Alexander Richter, R. Valas, M. Novotny, J. Yee-Greenbaum, J. D. Selengut, D. H. Haft, A. L. Halpern, R. S. Lasken, K. Nealson, R. Friedman, J. Craig Venter, Genomic insights to SAR86, an abundant and uncultivated marine bacterial lineage. ISME J. 6, 1186–1199 (2012).

46. K. Tamura, D. Peterson, N. Peterson, G. Stecher, M. Nei, S. Kumar, MEGA5: Molecular Evolutionary Genetics Analysis Using Maximum Likelihood, Evolutionary Distance, and Maximum Parsimony Methods. Mol. Biol. Evol. 28, 2731–2739 (2011).

47. S. F. Altschul, W. Gish, W. Miller, E. W. Myers, D. J. Lipman, Basic local alignment search tool. J. Mol. Biol. 215, 403–410 (1990).

48. E. Pruesse, J. Peplies, F. O. Glöckner, SINA: Accurate high-throughput multiple sequence alignment of ribosomal RNA genes. Bioinformatics. 28, 1823–1829 (2012).

49. S. M. S. M. Karst, R. H. Kirkegaard, M. Albertsen, Mmgenome: a Toolbox for Reproducible Genome Extraction From Metagenomes. bioRxiv, 059121 (2016).

50. R Core Team (2018) R: A language and environment for statistical computing R Foundation for Statistical Computing, Vienna, Austria (2018).

51. D. H. Parks, M. Imelfort, C. T. Skennerton, P. Hugenholtz, G. W. Tyson, CheckM : assessing the quality of microbial genomes recovered from isolates, single cells, and metagenomes. Genome Res. 25, 1043–1055 (2015).

52. B. J. Campbell, L. Yu, J. F. Heidelberg, D. L. Kirchman, Activity of abundant and rare bacteria in a coastal ocean. Proc. Natl. Acad. Sci. U. S. A. 108, 12776–12781 (2011).

53. T. Seemann, Prokka: Rapid prokaryotic genome annotation. Bioinformatics. 30, 2068–2069 (2014).

54. A. M. Eren, C. Esen, C. Quince, J. H. Vineis, H. G. Morrison, M. L. Sogin, T. O. Delmont, Anvi’o: an advanced analysis and visualization platform for ‘omics data. PeerJ, 1–29 (2015).

55. S. Kumar, G. Stecher, K. Tamura, MEGA7: Molecular Evolutionary Genetics Analysis version 7.0 for bigger datasets. Mol. Biol. Evol., 1–5 (2016).

56. R. C. Edgar, Search and clustering orders of magnitude faster than BLAST. Bioinformatics. 26, 2460–2461 (2010).

57. B. Bushnell, BBMap: A fast, accurate, splice-aware aligner Ernest Orlando Lawrence Berkeley National Laboratory, Berkeley, CA (US) (2014).

58. C. Quast, E. Pruesse, P. Yilmaz, J. Gerken, T. Schweer, P. Yarza, J. Peplies, F. O. Glöckner, The SILVA ribosomal RNA gene database project: improved data processing and web-based tools. Nucleic Acids Res. 41, D590–D596 (2013).

59. M. I. Love, S. Anders, W. Hu-, Differential analysis of count data – the DESeq2 package (2016).

60. J. Oksanen, F. G. Blanchet, R. Kindt, P. Legendre, P. R. Minchin, R. B. O’Hara, G. L. Simpson, P. Solymos, M. H. H. Stevens, H. Wagner, The vegan package. Community Ecol. Packag. 10, 631–637. (2007).

61. J. Juan, A. Armenteros, K. D. Tsirigos, C. K. Sønderby, T. N. Petersen, O. Winther, S. Brunak, G. Von Heijne, H. Nielsen, SignalP 5.0 improves signal peptide predictions using deep neural networks. Nat. Biotechnol. 37 (2019), doi:10.1038/s41587-019-0036-z.

62. N. Y. Yu, J. R. Wagner, M. R. Laird, G. Melli, S. Rey, R. Lo, P. Dao, S. C. Sahinalp, M. Ester, L. J. Foster, F. S. L. Brinkman, PSORTb 3.0: improved protein subcellular localization prediction with refined localization subcategories and predictive capabilities for all prokaryotes. Bioinformatics. 26, 1608–1615 (2010).

63. S. Ishii, S. Suzuki, A. Tenney, K. H. Nealson, O. Bretschger, Comparative metatranscriptomics reveals extracellular electron transfer pathways conferring microbial adaptivity to surface redox potential changes. ISME J. (2018), doi:10.1038/s41396-018-0238-2.

64. N. M. De Almeida, S. Neumann, R. J. Mesman, C. Ferousi, J. T. Keltjens, M. S. M. Jetten, B. Kartal, L. Van Niftrik, Immunogold Localization of Key Metabolic Enzymes in the Anammoxosome and on the Tubule-Like Structures of Kuenenia stuttgartiensis. J. Bacteriol. 197, 2432–2441 (2015).

65. B. Kartal, N. M. De Almeida, W. J. Maalcke, H. J. M. O. Den Camp, M. S. M. Jetten, J. T. Keltjens, N. M. De Almeida, W. J. Maalcke, H. J. M. Op den Camp, M. S. M. Jetten, J. T. Keltjens, How to make a living from anaerobic ammonium oxidation. FEMS Microbiol. Rev. 37, 428–461 (2013).

66. N. M. De Almeida, H. J. C. T. Wessels, R. M. De Graaf, C. Ferousi, M. S. M. Jetten, J. T. Keltjens, B. Kartal, Membrane-bound electron transport systems of an anammox bacterium : A complexome analysis. Biochim. Biophys. Acta - Bioenerg. 1857, 1694–1704 (2016).

67. J. Kostera, J. McGarry, A. A. Pacheco, Enzymatic interconversion of ammonia and nitrite: The right tool for the job. Biochemistry. 49, 8546–8553 (2010).

68. Z. He, J. Kan, Y. Wang, Y. Huang, F. Mansfeld, K. H. Nealson, Electricity Production Coupled to Ammonium in a Microbial Fuel Cell. Environ. Sci. Technol. 43, 3391–3397 (2009).

69. B. Qu, B. Fan, S. Zhu, Y. Zheng, Anaerobic ammonium oxidation with an anode as the electron acceptor. Environ. Microbiol. Rep. 6, 100–105 (2014).

70. A. Vilajeliu-Pons, C. Koch, M. D. Balaguer, J. Colprim, F. Harnisch, S. Puig, Microbial electricity driven anoxic ammonium removal. Water Res. 130, 168–175 (2018).

71. G. Zhan, D. Li, Y. Tao, X. Zhu, Ammonia as carbon-free substrate for hydrogen production in bioelectrochemical systems. Int. J. Hydrogen Energy. 39, 11854–11859 (2014).

72. G. Zhan, L. Zhang, D. Li, W. Su, Y. Tao, J. Qian, Autotrophic nitrogen removal from ammonium at low applied voltage in a single-compartment microbial electrolysis cell. Bioresour. Technol. 116, 271–277 (2012).

73. G. Zhan, L. Zhang, Y. Tao, Y. Wang, X. Zhu, Anodic ammonia oxidation to nitrogen gas catalyzed by mixed biofilms in bioelectrochemical systems. Electrochim. Acta. 135, 345–350 (2014).

74. M. Ruiz-Urigüen, W. Shuai, P. R. Jaffé, Electrode Colonization by the Feammox Bacterium Acidimicrobiaceae sp. Strain A6. *Appl. Environ*. Microbiol. 84, e02029–18 (2018).

75. M. Ruiz-Urigüen, D. Steingart, P. R. Jaffé, Oxidation of ammonium by Feammox Acidimicrobiaceae sp. A6 in anaerobic microbial electrolysis cells. Environ. Sci. Water Res. Technol. 5, 1582–1592 (2019).

76. B. Kartal, J. T. Keltjens, Anammox Biochemistry : a Tale of Heme c Proteins. Trends Biochem. Sci. 41, 998–1011 (2016).

77. A. Dietl, C. Ferousi, W. J. Maalcke, A. Menzel, S. de Vries, J. T. Keltjens, M. S. M. Jetten, B. Kartal, T. R. M. Barends, The inner workings of the hydrazine synthase multiprotein complex. Nature. 527, 394–397 (2015).

78. C. Ferousi, S. Lindhoud, F. Baymann, E. R. Hester, J. Reimann, B. Kartal, Discovery of a functional, contracted heme-binding motif within a multiheme cytochrome. J. Biol. Chem. (2019), doi:10.1074/jbc.RA119.010568.

79. M. Akram, J. Reimann, A. Dietl, A. Menzel, W. Versantvoort, M. S. M. Jetten, T. R. M. Barends, C. X. S. Group, A nitric oxide-binding heterodimeric cytochrome c complex from the anammox bacterium Kuenenia stuttgartiensis binds to hydrazine synthase. J. Biol. Chem. (2019), doi:10.1074/jbc.RA119.008788.

80. M. Akram, A. Dietl, U. Mersdorf, S. Prinz, W. Maalcke, J. Keltjens, C. Ferousi, N. M. de Almeida, J. Reimann, B. Kartal, M. S. M. Jetten, K. Parey, T. R. M. Barends, A 192-heme electron transfer network in the hydrazine dehydrogenase complex. Sci. Adv. 5, eaav4310 (2019).

81. L. Shi, H. Dong, G. Reguera, H. Beyenal, A. Lu, J. Liu, H. Yu, J. K. Fredrickson, Extracellular electron transfer mechanisms between microorganisms and minerals. Nat. Publ. Gr. 14, 651–662 (2016).

82. J. S. Gescher, C. D. Cordova, A. M. Spormann, Dissimilatory iron reduction in Escherichia coli: identification of CymA of Shewanella oneidensis and NapC of E. coli as ferric reductases. Mol. Microbiol. 68, 706–719 (2008).

83. C. R. Beckwith, M. J. Edwards, M. Lawes, L. Shi, J. N. Butt, D. J. Richardson, T. A. Clarke, Characterization of MtoD from Sideroxydans lithotrophicus: A cytochrome c electron shuttle used in lithoautotrophic growth. Front. Microbiol. 6, 332 (2015).

84. S. Luo, W. Guo, K. H. Nealson, X. Feng, Z. He, 13C Pathway Analysis for the Role of Formate in Electricity Generation by Shewanella Oneidensis MR-1 Using Lactate in Microbial Fuel Cells. Sci. Rep. 6, 1–8 (2016).

85. A. L. Kane, E. D. Brutinel, H. Joo, R. Maysonet, C. M. Vandrisse, N. J. Kotloski, J. A. Gralnick, Formate Metabolism in Shewanella oneidensis Generates Proton Motive Force and Prevents Growth without an Electron Acceptor. J. Bacteriol. 198, 1337–1346 (2016).

86. F. Kracke, I. Vassilev, J. O. Krömer, Microbial electron transport and energy conservation – the foundation for optimizing bioelectrochemical systems. Front. Microbiol. 6, 1–18 (2015).

87. Y. Liu, Z. Wang, J. Liu, C. Levar, M. J. Edwards, J. T. Babauta, D. W. Kennedy, Z. Shi, H. Beyenal, D. R. Bond, T. A. Clarke, J. N. Butt, D. J. Richardson, K. M. Rosso, J. M. Zachara, J. K. Fredrickson, L. Shi, A trans-outer membrane porin-cytochrome protein complex for extracellular electron transfer by Geobacter sulfurreducens PCA. Environ. Microbiol. Rep. 6, 776–785 (2014).

88. L. Shi, J. K. Fredrickson, J. M. Zachara, Genomic analyses of bacterial porin-cytochrome gene clusters. Front. Microbiol. 5, 1–10 (2014).

89. J. M. Dantas, M. A. Silva, D. Pantoja-uceda, D. L. Turner, M. Bruix, C. A. Salgueiro, Solution structure and dynamics of the outer membrane cytochrome OmcF from Geobacter sulfurreducens. BBA - Bioenerg. 1858, 733–741 (2017).

90. T. Ikeda, T. Ochiai, S. Morita, A. Nishiyama, E. Yamada, H. Arai, M. Ishii, Y. Igarashi, Anabolic five subunit-type pyruvate:ferredoxin oxidoreductase from Hydrogenobacter thermophilus TK-6. Biochem Biophys Res Commun. 340, 76–82 (2006).

91. R. H. H. Van Den Heuvel, B. Curti, M. A. Vanoni, A. Mattevi, Glutamate synthase : a fascinating pathway from L-glutamine to L-glutamate. Cell Mol Life Sci. 61, 669–681 (2004).

92. P. Y. Chen, B. Li, C. L. Drennan, J. Sean, P. Y. Chen, B. Li, C. L. Drennan, S. J. Elliott, A Reverse TCA Cycle 2-Oxoacid : Ferredoxin Oxidoreductase that Makes C-C Bonds from CO 2 A Reverse TCA Cycle 2-Oxoacid : Ferredoxin Oxidoreductase that Makes C-C Bonds from CO 2. Joule, 1–17 (2018).

93. K. A. Weber, L. A. Achenbach, J. D. Coates, Microorganisms pumping iron: anaerobic microbial iron oxidation and reduction. Nat Rev Microbiol. 4, 752–764 (2006).

94. N. Noinaj, M. Guillier, T. J. Barnard, S. K. Buchanan, TonB-Dependent Transporters : Regulation, Structure, and Function. Annu Rev Microbiol. 64, 43–60 (2010).

95. K. Mosbahi, M. Wojnowska, A. Albalat, D. Walker, Bacterial iron acquisition mediated by outer membrane translocation and cleavage of a host protein. Proc Natl Acad Sci U S A. 115, 6840–6845 (2018).

